# Architecture of the UBR4 complex, a giant E4 ligase central to eukaryotic protein quality control

**DOI:** 10.1101/2024.12.18.629163

**Authors:** Daniel B Grabarczyk, Julian F Ehrmann, Paul Murphy, Robert Kurzbauer, Lillie E Bell, Luiza Deszcz, Jana Neuhold, Alexander Schleiffer, Alexandra Shulkina, Gijs A Versteeg, Anton Meinhart, Eszter Zavodszky, Ramanujan S Hegde, Tim Clausen

**Affiliations:** Research Institute of Molecular Pathology, Vienna BioCenter (VBC), Vienna, Austria; Vienna BioCenter PhD Program, Doctoral School of the University of Vienna and Medical University of Vienna, Vienna BioCenter (VBC), Vienna, Austria; Department of Biological Chemistry and Molecular Pharmacology, Harvard Medical School, Boston, MA, USA; Vienna Biocenter Core Facilities, Vienna BioCenter, Vienna, Austria; Max Perutz Labs, Vienna Biocenter (VBC), Vienna, Austria; University of Vienna, Center for Molecular Biology, Department of Microbiology, Immunobiology, and Genetics, Vienna, Austria; MRC Laboratory of Molecular Biology, Cambridge, UK; Medical University of Vienna, Vienna, Austria

**Author notes:** To whom correspondence should be addressed: DBG, TC.

## Abstract

Eukaryotic cells have evolved sophisticated quality control mechanisms to eliminate aggregation-prone proteins that compromise cellular health. Central to this defense is the ubiquitin-proteasome system, where UBR4 acts as essential E4 ubiquitin ligase, amplifying degradation marks on defective proteins. Our cryo-EM analysis of UBR4 in complex with its cofactors KCMF1 and CALM1 reveals a massive 1.3 MDa ring structure, featuring a central substrate-binding arena and flexibly attached catalytic units. Structural data illustrate how UBR4 binds substrate and extends K48-specific ubiquitin chains. Importantly, efficient substrate targeting depends on both pre-ubiquitination and specific N-degrons, with KCMF1 acting as key substrate filter. Furthermore, we show that the architecture of the E4 megacomplex is conserved across eukaryotes but with species specific adaptations, allowing UBR4 to perform its precisely tuned quality-control function in diverse cellular environments.

## Introduction

Protein function depends on proper folding, complex assembly, and precise localization within the cell. When any of these processes fail, misfolded proteins can build up, forming harmful aggregates that compromise cell health (*1*). To safeguard against such proteotoxic damage, all cells employ sophisticated protein quality control (PQC) mechanisms, with the ubiquitin-proteasome system (UPS) acting as a key clean-up pathway that tags defective proteins with ubiquitin molecules for rapid destruction (*2, 3*).

Specificity of ubiquitination is determined by E3 ubiquitin ligases (*4*). The best characterized E3 ligases recognize distinct sequence or structural motifs (*5*), allowing the tightly controlled removal of specific target proteins, as required, for example, for cell cycle regulation (*6*). In contrast, E3 ligases involved in PQC must be able to target diverse proteins that are present in a non-native state, but otherwise do not share specific molecular features (*7*). Accurate function of PQC ligases is ensured by intricate binding mechanisms involving various receptor sites (*8, 9*) and by adaptor proteins that specifically recognize defective substrates (*10, 11*), thus preventing the mislabeling of functional proteins. An additional security measure is provided by UPS enzymes that directly modify the degradation signal. While deubiquitinases remove ubiquitin tags and stabilize specific targets (*12*), certain E3 ligases enhance degradation by amplifying the ubiquitination signal. The latter class of enzymes, known as E4 ligases (*13*), not only extend poly-ubiquitin chains but also generate branched chains, which are particularly strong proteasomal degradation signals (*14–16*). Given their critical role in maintaining cellular homeostasis, it is not surprising that ubiquitin ligases have been linked to diverse diseases including neurodegeneration, myopathies and cancer (*17–19*). Aneuploidy, for example, disrupts subunit stoichiometry, generates orphan proteins, and imposes proteotoxic stress, highlighting the importance of PQC ligases as potential therapeutic targets (*20, 21*).

Among the >600 ubiquitin ligases in human cells (*22*), UBR4 plays a particularly important PQC role, being an essential and conserved E3/E4 ligase found across eukaryotes (*23*). UBR4 is critical for maintaining proteostasis in long-lived cells with high metabolic demands, such as neurons and muscle cells, where proteotoxic stress is particularly harmful (*24–27*). Recent evidence has shown that UBR4 recognizes multiple substrate features, including mitochondrial targeting sequences (MTS) of mislocalized mitochondrial precursors, orphan proteins resulting from unassembled complexes, and aggregation-prone proteins (*28–31*). Additionally, UBR4 plays a key role in autophagic processes, bridging proteasomal and lysosomal degradation pathways to ensure efficient clearance of aberrant proteins (*27*). The diverse functions of UBR4 underscores its role as a central hub in the PQC network, capable of integrating multiple stress response pathways and teaming up with various ubiquitin ligases. At the same time, UBR4 activity seems to rely on two specific cofactors: the calcium-binding protein calmodulin (CALM1) and KCMF1 (*25, 28, 29, 32*). Although KCMF1 has been reported to function as an E3 ligase (*29, 33*), its precise role within the UBR4 complex remains unclear, particularly as it is absent in many organisms. To elucidate how these proteins interact to assemble an efficient yet tightly regulated E4 ligase, we reconstituted the UBR4/KCMF1/CALM1 complex and examined its structure and function. Our findings reveal that UBR4 acts as a K48-linkage-specific E4 ligase, employing KCMF1 not as a partner ubiquitin ligase but as a tightly bound adaptor protein, targeting pre-ubiquitinated with a specific N-degron or MTS. The 1.3 MDa hexameric complex adopts a ring-like architecture, comprising a central substrate-binding arena that is read out by a flexibly attached catalytic unit. We further demonstrate that the overall structure and mechanism of UBR4 complexes are conserved across eukaryotes, highlighting its role as a central PQC factor.

## Results

### Structure of the human UBR4 complex reveals a ring-shaped ubiquitination arena

To investigate how UBR4 and its KCMF1 and CALM1 cofactors target proteins for degradation, we reconstituted the human E4 complex with recombinant proteins expressed in insect cells. We either purified the three proteins separately before mixing them together or co-expressed them to purify the ternary complex via a tag on UBR4. When we assessed the composition of the E4 complex by mass photometry, we observed that the co-expressed complex contained UBR4, KCMF1, and CALM1, and had a molecular weight consistent with a dimer of heterotrimers (**Fig. S1A**). In contrast, UBR4 isolated separately was monomeric and did not associate with KCMF1 and calmodulin (**Fig. S1B**). Moreover, monomeric UBR4 showed a tendency to aggregate and become degraded, while the co-expressed complex was stable.

We next determined the structure of the human UBR4 complex (HsUBR4_2_/KCMF1_2_/CALM1_2_) using cryo-electron microscopy (cryo-EM). Through focused classification and refinement, we obtained density maps encompassing all folded sections of each component (**Fig. 1A, Fig. S2, Fig. S3**), except the flexible C-terminal domain containing the hemi-RING E3 module (*34*). Overall, two UBR4 molecules assemble a through two distant dimerization interfaces, resulting in the formation of a large molecular ring (**Fig. 1A**). One dimer interface is formed by the N-terminal domain, involving extensive contacts between adjacent Armadillo repeats (**Fig. 1B**). The second interface, which is much larger, is formed by Armadillo repeats in the C-terminal portion and laterally aligned helices sealing this contact. This interface is further appended by the partner proteins CALM1 and KCMF1, as previously predicted for the trimeric complex (*28*). The C-terminal helix of KCMF1 acts as a pin, inserting into a hole in the helical scaffold of UBR4. Here, it forms one helix of the Armadillo repeat and is covered by a small lid insertion within the region that also mediates CALM1 binding. The intricate KCMF1-UBR4 binding mode explains why UBR4 is unstable when it is not co-expressed with its cofactor: both the Armadillo repeat and this lid structure would be unlikely to fold properly without KCMF1. To confirm the importance of this interaction, we co-expressed UBR4 with a KCMF1 variant where hydrophobic residues in the C-terminal helix were mutated (L318A/F319A/V320S). The resultant UBR4 protein was monomeric and unstable (**Fig. S1C**), as predicted by the cryo-EM model.

**Fig. 1.**
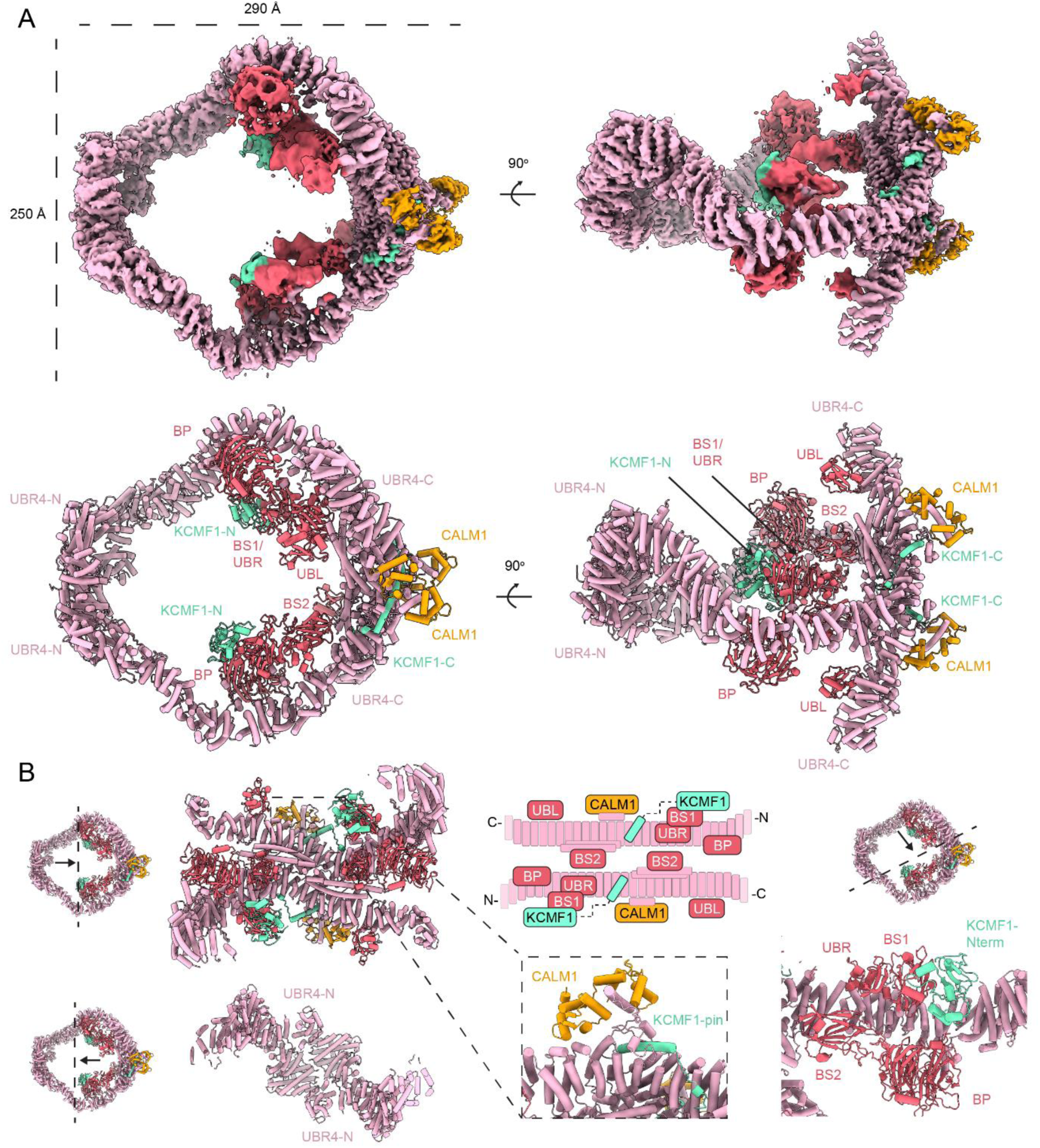
Architecture of the human UBR4 complex. **(A)** Composite cryo-EM density map and model colored by protein component and domain as indicated. **(B)** Detailed views of structural features of the complex.

CALM1 often mediates calcium regulation of target proteins and is bound in our structure in a manner typical of other calmodulin-interacting proteins with a long hydrophobic helix docking into the cleft between the two lobes of CALM1 (**Fig. 1B**) (*35, 36*). The C-lobe of CALM1 adopts a calcium-bound conformation (**Fig. S1D**), suggesting that calmodulin binding to UBR4 may depend on the availability of calcium and, accordingly, we observed a significant CALM1-free UBR4 population in our cryo-EM dataset despite CALM1 overexpression (**Fig. S1E, Fig. S2**). Addition of the calcium-chelator EGTA did not affect the CALM1 occupancy in the structure, suggesting that once CALM1 is bound it remains stable rather than acting as a reversible regulator (**Fig. S2**).

Whereas the structural motifs forming the rigid scaffold of the E4 ring were well defined by EM density, the functional domains lining the inner cavity or extending into the periphery exhibited high flexibility, as reflected by poorer local resolution (**Fig. S1E, Fig. S2**). The most prominent intrusion into the E4 ring is formed by a beta-propeller (BP) and an attached multidomain structure comprising the UBR box and beta-sandwich 1 (BS1) domains of UBR4 together with the N-terminal zinc binding domains of KCMF1 (**Fig. 1B**). A second appendix protrudes from the C-terminal dimerization region, where a zinc finger domain positions another beta-sandwich domain (BS2) with two associated zinc fingers near the central cavity. Finally, an extension projects outwards from the UBR4 ring, and is formed by the C-terminal Armadillo repeats. This structure houses a ubiquitin-like (UBL) domain and a flexibly attached hemi-RING module, which was unresolved in the EM map.

### Ubiquitin K48 chain extension within the UBR4 ring

To investigate how the distinct functional motifs of the 1.3 MDa UBR4 complex contribute to ubiquitination activity, we developed an E4 assay monitoring the conjugation of the two ubiquitin variants Ub* and Ub-K0 (**Fig. 2A**). Ub* (ubiquitin-ΔGG) lacks the C-terminal residues required for ubiquitin transfer, while Ub-K0 has all lysine residues mutated to arginine. This setup enabled us to follow the rate of a single ubiquitination event where Ub-K0 is transferred onto Ub*, and then no further reaction can occur. As anticipated, a Ub-K0-Ub* band was observed only in the presence of both ubiquitin variants and UBR4 (**Fig. 2A**), offering a clear and specific readout of E4 activity, ideal for mutational analyses and systematic substrate screens.

**Fig. 2.**
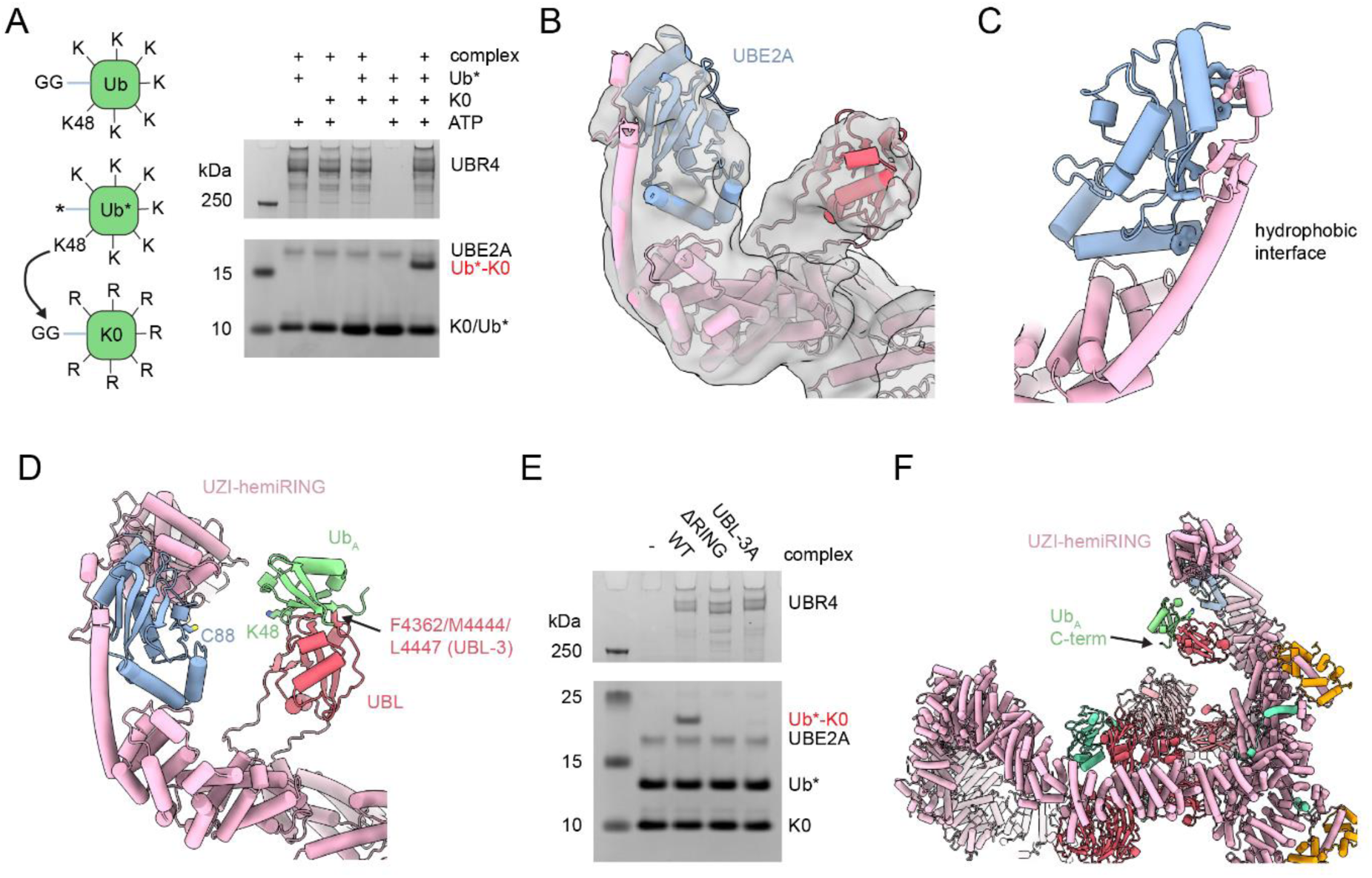
Mechanism of ubiquitin chain extension by the UBR4 complex. **(A)** E4 ligase assay showing formation of K0-Ub-Ub* di-ubiquitin catalysed by 200 nM human UBR4 complex when all components of the assay are added for 45 minutes at 37°C. **(B)** Cryo-EM density map and model of the structure of human UBR4 complex obtained in the presence of UBE2A. **(C)** Detailed view of the interaction highlighting the two-sided interface with hydrophobic interactions. **(D)** Model of the E4 transfer state using our structure combined with a crystal structure of the hemiRING-UBE2A complex (PDB 8BTL) and an AlphaFold3 model of the UBL-Ub interaction. **(E)** Zoomed out image of the model showing the position of the C-terminal tail of Ub relative to the UBR4 arena. **(F)** E4 ligase assay with 100 nM of the indicated variant complexes for 45 minutes at 37°C.

We previously observed that UBR4 only ubiquitinates orphan proteins which have already been ubiquitinated by other E3 ligases (*28*). This E4 activity was dependent on the UBL domain which was predicted by AlphaFold3 (*37*) to bind ubiquitin and position lysine K48 towards the E2∼Ub conjugate bound to the hemi-RING, thereby promoting formation of K48-linked chains on pre-ubiquitinated substrates (*28*). To further human UBR4 complex with its cognate E2, UBE2A, and performed cryo-EM analysis (**Fig. S4, Table S1**). We now observed additional density for UBE2A bound to the end of the helical UBR4 scaffold (**Fig. 2B**). As predicted by AlphaFold3 (*28*), the E2 is bound in a backside configuration, stabilized by a specific beta-hairpin structure within an extended helix of the C-terminal protrusion. This interaction features an extensive hydrophobic interface that spans two sides of UBE2A (**Fig. 2C**), while leaving the canonical E3/E4 binding site accessible for interaction with the hemi-RING domain of UBR4. The stable interface precisely positions the E2 at the narrow constriction site of the substrate arena, bringing it into close proximity with the UBL domain.

With regards to the catalytic module, we did not observe the hemi-RING domain in our cryoEM map due to the inherent flexibility of the C-terminal extension. We thus modelled the ubiquitin transfer state by aligning the UBE2A-UBR4(hemi-RING) crystal structure (*34*) onto the UBE2A in our cryo-EM model (**Fig. 2D**), along with the AlphaFold3 prediction of the UBL-Ub interaction. In this structure guided model, UBE2A and the UBL domain were well positioned to orient the acceptor ubiquitin towards the hemi-RING, with the K48 residue protruding towards the E2∼Ub thioester to enable ubiquitin chain extension. A triple mutation in the ubiquitin receptor site of the UBL domain (UBL-3A) or deletion of the hemi-RING domain (ΔRING) abrogated ubiquitin conjugation, confirming the central role of the UBL domain in the E4 mechanism of UBR4 (**Fig. 2E**). For native substrates, the ubiquitin captured by the UBL domain would be linked via its C-terminus to a lysine residue on the target protein. Notably, the ubiquitin C-terminus points into the central UBR4 cavity, suggesting the presence of potential substrate recognition sites within this region (**Fig. 2F**).

Previous studies suggested that KCMF1 functions as an E3 ligase (*29, 33, 38*), and thus could potentially initiate substrate ubiquitination. However, these earlier results may reflect an indirect effect, as our findings indicate that the C-terminus of KCMF1 is essential for stabilizing UBR4. A closer analysis of KCMF1 revealed that it does not contain any known E3 domains (**Fig. S5A**). The N-terminal zinc binding domain (1–80) has similar topology to a RING domain but is a ZZ domain with high sequence similarity to these domains in p62 (*39, 40*), ZZZ3 (*41*) and p300 (*42*) which recognize specific N-terminal residues but have no E3 activity. The other zinc binding domain (80–160) is a Di19-Zn-binding (DZB) domain which has distinct topology to a RING domain. The only reported function of a DZB domain is binding mono-ADP-ribosylated histones (*43*). Consistent with these observations, when we tested the ubiquitination activity of isolated KCMF1 with a panel of human E2 enzymes, no activity was detected (**Fig. S5B**). Taken together, our data indicate that the human UBR4 complex operates as a bona fide E4 ligase, ready to collaborate with partner E3 ligases in the PQC network.

### Selection of Ub-marked proteins to be degraded

UBR4, with a mobile E4 module on top of a substrate binding arena, mirrors the architecture of giant ubiquitin ligases like BIRC6, HUWE1 and UBR5, all of which use an array of receptor domains to target diverse substrates (**Fig. S6A**) (*8, 9, 44–48*). To test whether UBR4 may also target distinct substrate classes, we first performed a co-IP/MS experiment in human cells to categorize potential target proteins (**Fig. 3A**). Consistent with earlier studies, we detected many mitochondrial proteins (*25, 29*). These have been reported to be recognized via their mitochondrial targeting sequence (MTS) and accordingly, we detected several unprocessed MTS peptides in our co-IP/MS data (**Fig. S6B**). The MTS-bearing proteins could be direct substrates of UBR4 as previously suggested or, in accordance with our E4 model, are only recognized because they have been pre-ubiquitinated by other E3 ligases. To distinguish between these two models, we performed a co-IP/MS experiment with a lysate that was treated with USP21, a non-specific deubiquitinase (DUB), to remove all ubiquitination marks. Comparison of the DUB-treated to the untreated sample revealed a striking depletion of mitochondrial proteins (**Fig. 3B**). These data suggest UBR4 acts as an E4 ligase for mislocalized pre-ubiquitinated mitochondrial proteins similarly to its role in targeting pre-ubiquitinated orphaned proteins for proteasomal degradation (*28*).

**Fig. 3.**
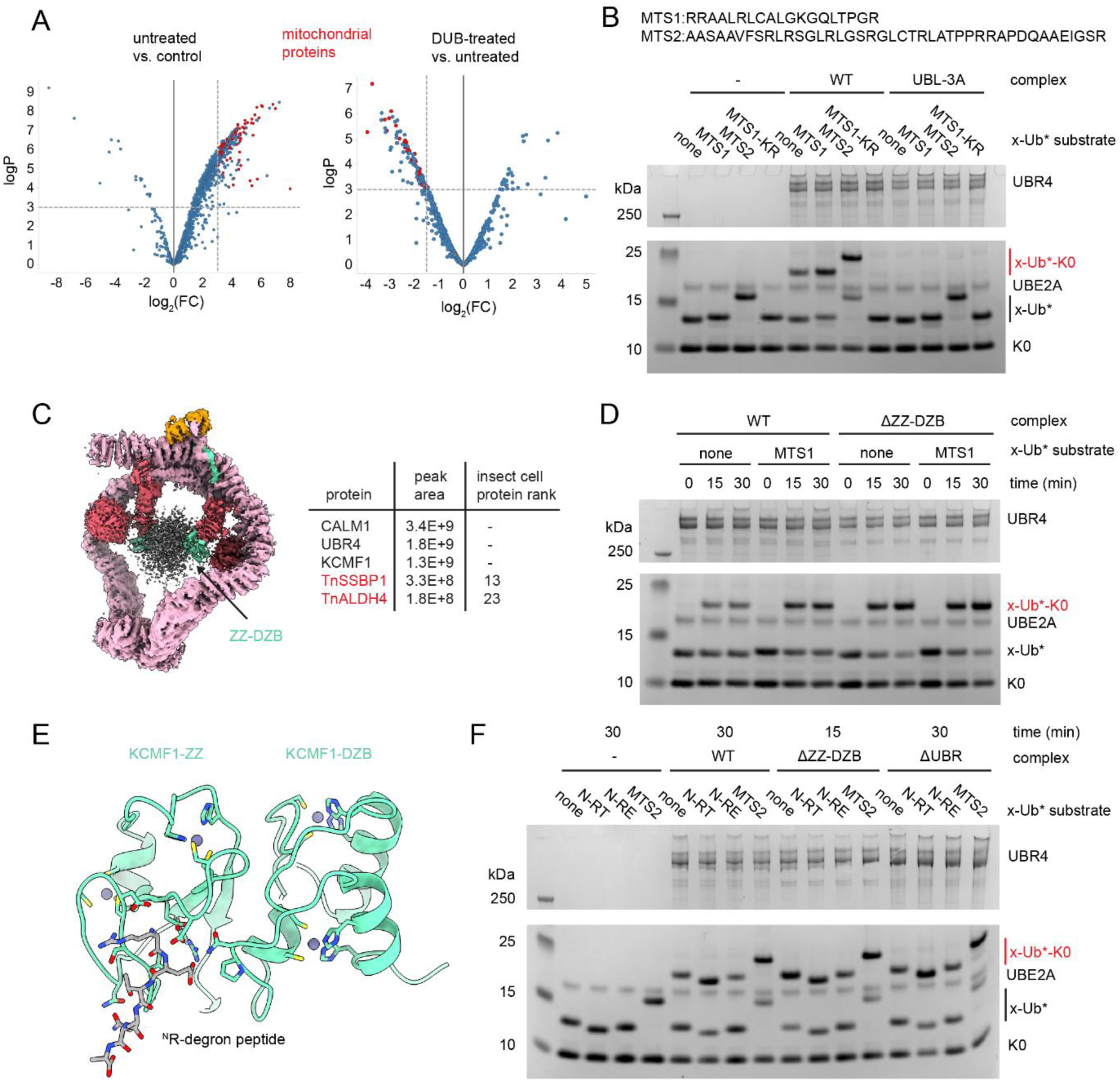
KCMF1 mediates selection of UBR4 substrates. **(A)** Co-IP/MS volcano plots comparing a UBR4 pull-down vs GFP control on the left and on the right UBR4 pull-down from lysates which were treated vs untreated with the DUB USP21. Mitochondrial proteins are highlighted in red. **(B)** E4 ligase assay with 200 nM of the indicated human UBR4 complexes for 30 minutes at 37°C with the indicated X-linker-Ub* or X-linker-Ub(K48R)* substrates. **(C)** On the left, the human UBR4 complex structure refined with a global mask showing fuzzy central substrate density. On the right, a table summarizing MS analysis of insect cell contaminants in the purified sample with mitochondrial proteins shown in red. **(D)** E4 ligase assay as in panel **B** with the indicated substrates, complexes and reaction times. **(E)** Alphafold3 model of the KCMF1 ZZ-DZB domains with an ^N^RE N-degron sequence. **(F)** E4 ligase assay as in panel **D** with different N-degron Ub* substrates. A shorter incubation time was used for the ZZ-DZB construct due to its stronger activity on all substrates as shown in panel D.

Despite the requirement for pre-ubiquitination for recognition, the enrichment of mitochondrial precursors among all ubiquitinated proteins suggests that UBR4 is able to recognize specific protein motifs. To test for a respective substrate filter, we adapted our single ubiquitination E4 assay by fusing candidate sequences, via a flexible linker, to the Ub* N-terminus (**Fig. 3C**). We then compared activity against our control (linker-Ub*) and model (MTS-linker-Ub*) substrates, using targeting sequences from two co-IP/MS hits, ACOT9 (MTS1) and TIMM50 (MTS2) (**Fig. 3C**). We observed stronger ubiquitination of the MTS fused substrates compared to linker-Ub* alone, reflecting the inherent specificity of UBR4 for mitochondrial precursors. To test whether we were measuring UBL-mediated E4 rather than direct E3 ligase activity, we used the UBL-3A complex variant which disrupts the ubiquitin interaction site, thus abolishing chain extension without compromising hemi-RING E3 activity. Importantly, this mutation fully abolished ubiquitination of our MTS1-Ub* and MTS2-Ub* model substrates (**Fig. 3B**). Moreover, introducing a K48R mutation into one of the MTS-Ub* model substrates, which should also prevent K48 chain formation, impaired product formation as well (**Fig. 3B**). Together, these data demonstrate that the human UBR4 complex is a substrate-specific E4 ligase.

When investigating how substrates are recognized, we noted that our cryo-EM reconstruction contained extra low-resolution density in the central cavity of the UBR4 ring (**Fig. 3C**). As the flexible C-terminal extension is too distant to account for this density, we considered a copurified protein, such as a substrate, that was captured in the inner cavity, providing direct insight into the mechanism of substrate recognition. To look for the captured target, we re-analyzed MS data of the purified complex (**Fig. 3C**). Among common protein contaminants, we observed a significant enrichment of single stranded DNA binding protein (SSBP1), a mitochondrial protein previously reported to bind to UBR4/KCMF1 in human cells (*25*). While the low resolution did not allow fitting of a molecular model into the density in the inner cavity, we observed a clear connection of this density to the ZZ-DZB domains of KCMF1 (**Fig. 3C**), consistent with previous observations that this domain is involved in binding mitochondrial proteins (*25*). To test whether the ZZ-DZB is the recognition site for MTS sequences, we reconstituted the UBR4_2_/KCMF1(Δ2-138)_2_/CALM1_2_ complex lacking the putative receptor site and tested the E4 activity against our MTS model substrates. For the mutated complex, ubiquitination of the MTS1-Ub* substrate was now indistinguishable from the linker-Ub* control, suggesting that substrate specificity has been lost (**Fig. 3D**). Surprisingly, the activity of ΔZZ-DZB complex was stronger than the WT complex towards both the MTS1-Ub* and Ub* substrates. This result suggests that the ZZ-DZB is not only a substrate receptor but also functions as a selectivity filter, restricting access to the site within the central cavity where substrates are subjected to E4 activity.

KCMF1 has been previously reported to recognize type I Arg N-degrons via its ZZ domain (*38*). Consistently, AlphaFold3 predicts a complex between the KCMF1 ZZ-domain and an ^N^R-peptide motif (**Fig. 3E**). Notably, our model substrate MTS1-Ub* has an N-degron-like ^N^RR-sequence, but the equally well targeted MTS2-Ub* does not, bearing an N-terminal ^N^AA-sequence. To explore potential N-degrons, we started with our control construct (^N^TA-linker-Ub*) and generated an ^N^RT-degron sequence (^N^RTA-linker-Ub*) as well as the ^N^RE-degron (^N^RETA-linker-Ub*) previously shown to bind the ZZ domain of p62 (*39, 40*). The ^N^RT-Ub* construct was ubiquitinated as strongly as the MTS2-Ub* construct (**Fig. 3F**), showing that the UBR4 complex specifically recognizes type I N-degrons. In contrast, the ^N^RE degron was targeted worse than the control sequence. The preference for the ^N^RT degron was dependent on the ZZ-DZB domain of KCMF1 suggesting that recognition of N-degrons is mechanistically similar to that of MTS sequences. Both the ^N^RT degron and the MTS sequences are characterized by their basicity and so it is possible that the UBR4 complex disfavors binding of negative charges, as present in the ^N^RE sequence. Of note, UBR4 was initially identified as an N-degron ubiquitin ligase due to its UBR domain, a motif associated with recognizing N-degrons. Yet, our data indicate that the UBR domain is not essential for recognition of substrates tested in this study, as its deletion did not impair the ubiquitination of MTS-Ub* or ^N^R-Ub* (**Fig. 3F**). These findings emphasize the role of the ZZ-DZB as substrate receptor within the UBR4 complex.

Furthermore, our results help clarify why earlier studies suggested KCMF1 might function as an E3 ligase together with UBR4, as mutations in the ZZ-DZB domain disrupt substrate recognition. To clearly rule out an E3 function for KCMF1 within the UBR4 complex, we analyzed ubiquitination in the presence and absence of a UBE2D family E2, the reported partner of KCMF1 (*33*), using wild-type ubiquitin capable of chain formation and the MTS1-Ub* substrate. UBE2D2, when tested without UBE2A, was unable to support any E3 activity on this substrate. Furthermore, its addition did not alter the activity catalyzed by UBE2A or its dependence on the K48 residue of the Ub* substrate moiety, despite there being an exposed lysine within the MTS (**Fig. S5C**).

In conclusion, we show that the ZZ-DZB is not an E3 module but a specialized substrate receptor within the UBR4 complex. The low specificity of this domain fits well with the general PQC role of UBR4, targeting a broad range of MTS and N-degron sequences.

### Evolutionary conservation of the UBR4 ubiquitination arena

UBR4 has been shown to play essential roles in various eukaryotes, including *Arabidopsis thaliana* (*49, 50*), *Drosophila melanogaster* (*51, 52*), and *Caenorhabditis elegans* (*53*). Notably, UBR4 from nematodes and plants has not been associated with KCMF1, a protein that is absent outside of metazoans and considerably larger in nematodes. This raises the question of whether the structurally and functionally characterized human UBR4 complex reflects the universal mechanisms underlying the role of UBR4 in eukaryotic protein quality control.

To address this question, we first focused on *C. elegans* UBR4, which has been linked to orphan protein degradation (*53*) and for which we could identify a potential KCMF1 orthologue, ZK652.6. We co-expressed CeUBR4, CeKCMF1, and CeCALM, purified the resulting complex, and compared it to CeUBR4 alone. Like the human proteins, CeUBR4 was monomeric when expressed alone, while the co-expressed complex exhibited a molecular weight of 1.1 MDa consistent with a dimer of trimers (**Fig. S7A**). However, MS analysis revealed that CeCALM was not retained in the purified complex. Activity assays demonstrated that the CeUBR4 complex (UBR4_2_/KCMF1_2_) exhibited strong selectivity for MTS-Ub* substrates over Ub* alone but showed reduced activity on the ^N^RT-Ub* degron substrate (**Fig. 4A**). Overall, these findings indicate functional conservation between human and *C. elegans* UBR4, while also highlighting species specific adaptations in substrate selectivity and complex composition.

**Fig. 4.**
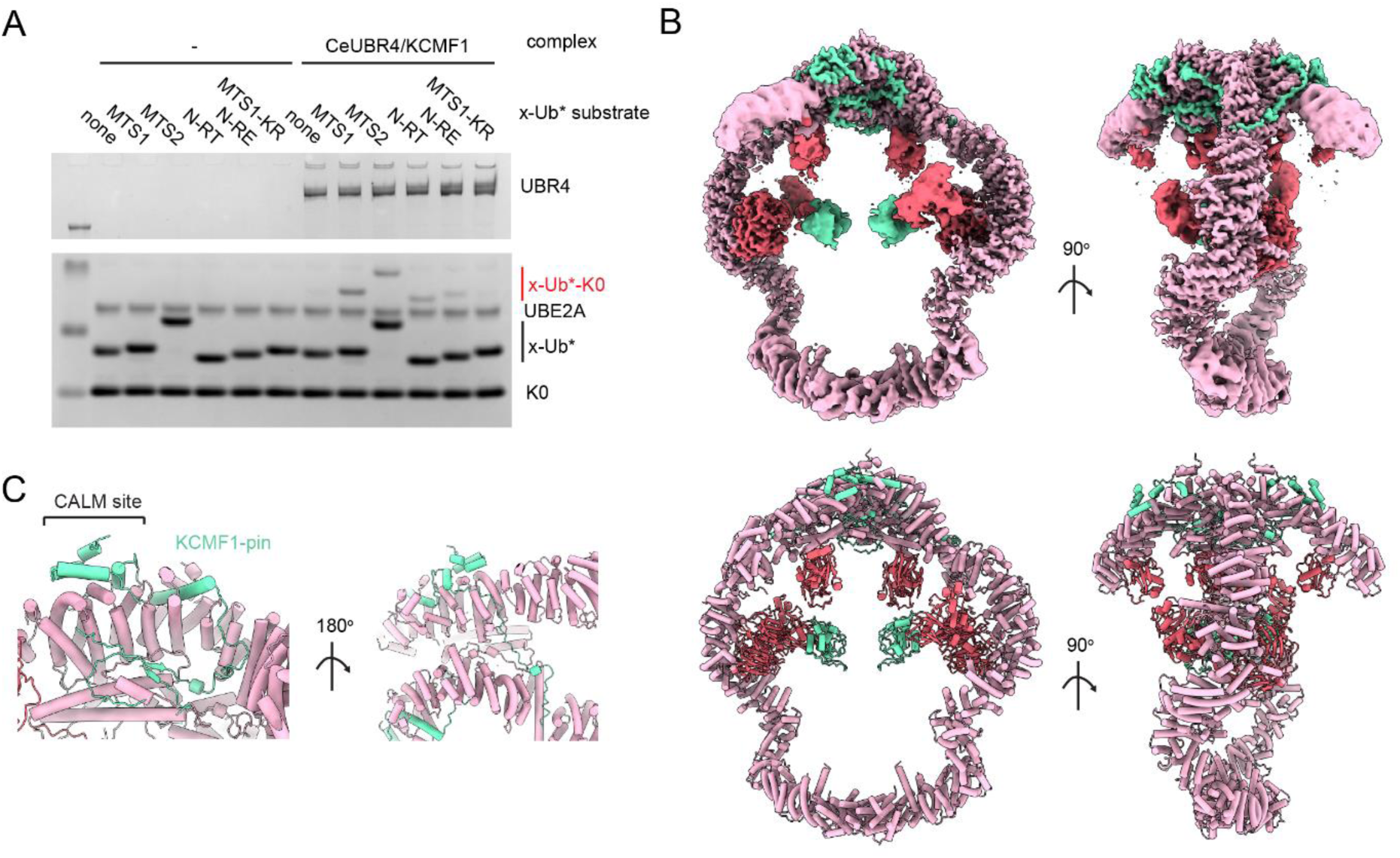
Structure of the CeUBR4 complex. **(A)** E4 ligase assay using 200 nM CeUBR4 complex and the indicated Ub* substrates for 45 minutes at 37°C. **(B)** Cryo-EM model and map of the CeUBR4 complex with different orientations. Domains and subunits are coloured as in Fig. 1 **(C)** Detailed view of the extended interface of CeKCMF1 with CeUBR4.

To explore the underlying structural differences, we determined the cryo-EM structure of the CeUBR4 complex (**Fig. 4B, Table S1, Fig. S8, Fig. S9**). While the dimeric complex retains the ring-shaped organization of the human counterpart, it exhibits significant structural adaptations. Notably, CALM is absent and replaced by insertions in CeKCMF1 (**Fig. 4C**). The conserved C-terminal helical pin of KCMF1 interacts with UBR4, but this interaction is further stabilized by a mixed beta-sheet that connects KCMF1 with both UBR4 protomers, effectively gluing the dimer together. Additionally, two helices from KCMF1 extend from the mixed beta-sheet to engage the same interface on UBR4 that CALM1 interacts with in the human structure. The necessity of occupying this interface in the absence of CALM1 highlights its critical role in maintaining the structural integrity of the E4 complex. Supporting a structural rather than regulatory function, deletion of the CALM1-interacting helix in HsUBR4 had no effect on its E4 activity (**Fig. S7B**). Beyond this interface, an extended C-terminal peptide of KCMF1 wraps tightly around the back of the UBR4 dimer, aligning the UBR4 protomers from the opposite side. The modified CeUBR4/CeKCMF1 interface alters the overall conformation of the UBR4 ring, a distortion further reinforced by a distinct dimerization interface at the opposite end, involving unique N-terminal domains (**Fig. 4B**). Notably, the relative positions and structures of key functional units—including the UBL, BP, and BS2 domains, the UBR-BS1-ZZ-RING composite structure, and the flexible C-terminal extension housing the E4 active site remain consistent with the human UBR4 complex, emphasizing the evolutionary conservation of its core architecture.

While the CeUBR4 complex retains E4 activity on mitochondrial precursors, its architecture shows pronounced differences to its human counterpart. This divergence prompted us to investigate the ancestral design of the UBR4 core further. We turned to *Arabidopsis thaliana* UBR4 (also known as BIG), which diverged from metazoan UBR4 early in eukaryotic evolution. Unlike metazoan UBR4, plants lack a KCMF1 orthologue, raising questions about how UBR4 architecture and activity are maintained. We expressed and purified AtUBR4 from insect cells and observed that the protein was homodimeric (**Fig. S7C**) and possessed detectable but weak substrate- and K48-specific E4 ligase activity, despite the lack of the KCMF1 co-factor (**Fig. 5A**). Furthermore, in negative stain analysis, we saw a ring structure with similar dimensions to the human UBR4 complex (**Fig. 5B**). We performed MS analysis to identify co-purified components of the insect cell UBR4 complex and found strong enrichment of calmodulin but no detectable KCMF1 (**Fig. 5C, Table S3**). Consistently, the CALM1-interacting helix of HsUBR4 is well conserved in AtUBR4 (**Fig. 5D**). Considering the apparent structural and functional similarity between plant and human UBR4, we hypothesized that a so-far undiscovered KCMF1 orthologue should stabilize AtUBR4. We identified a candidate protein, drought-induced protein 19 (DI19), which shows strong homology to HsKCMF1 in the C-terminal pin region (**Fig. 5E**), has related biological functions to AtUBR4 (*49, 54*) and was one of the top hits along with AtUBR4 in a proximity labelling experiment using an N-degron as bait (*50*). Using AlphaFold3, we were able to predict the AtUBR4_2_/DI19_2_/CALM_2_ complex with high confidence (**Fig. 5E**), confirming the identity of the plant KCMF1. Thus, the structural core of the UBR4 complex has been maintained throughout eukaryotic evolution.

**Fig. 5.**
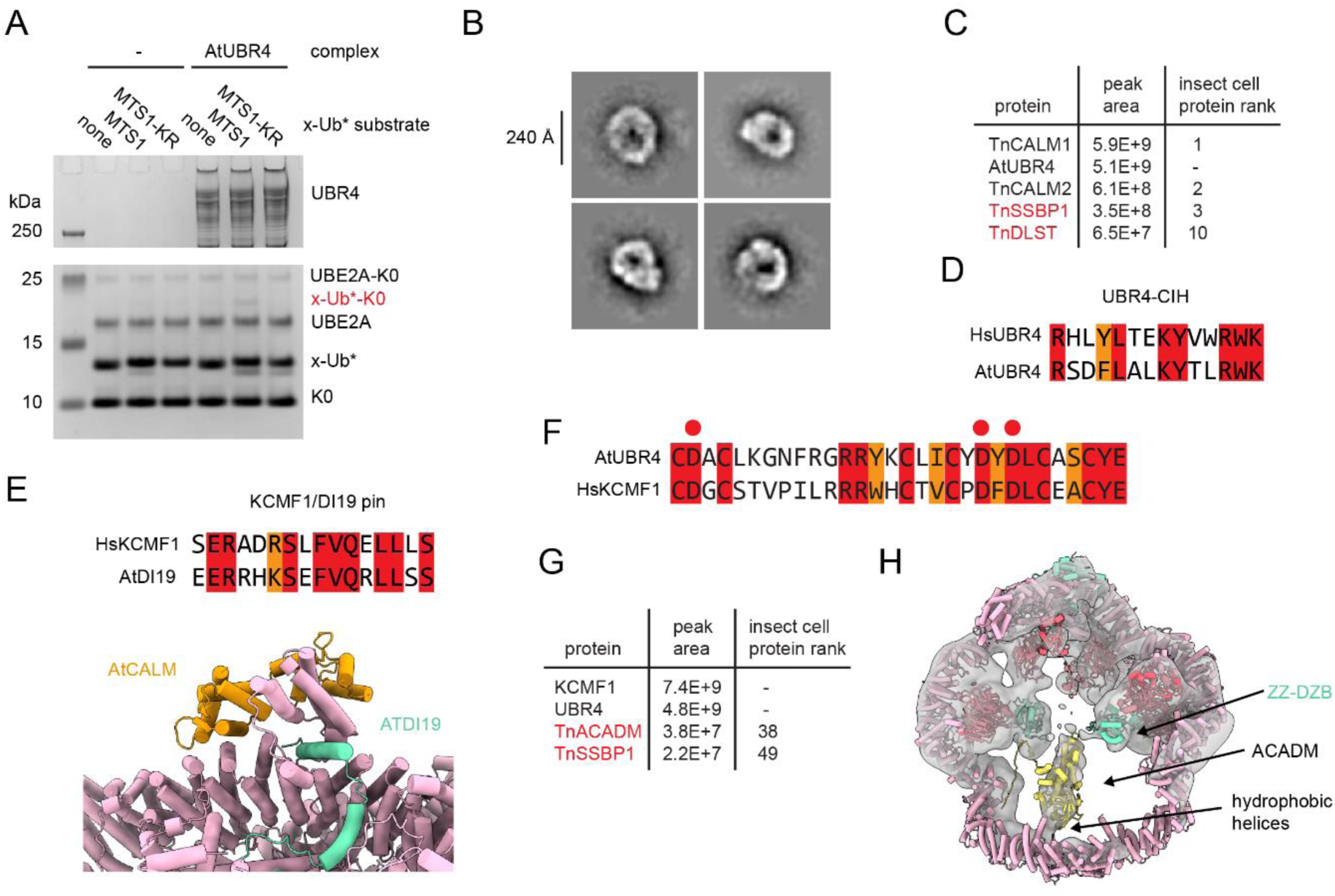
Evolutionary conservation and adaptations of the UBR4 complex. **(A)** E4 ligase assay with 400 nM AtUBR4 for 90 minutes at 37°C. **(B)** Negative stain 2D classes of AtUBR4. **(C)** Summary of MS data of purified AtUBR4 showing ranking of insect cell contaminants with mitochondrial proteins in red. **(D)** Conservation of the UBR4 CALM-interacting helix between HsUBR4 and AtUBR4. **(E)** Conservation of the KCMF1 pin between HsKCMF1 and AtDI19 and an Alphafold3 model of the AtUBR4_2_/AtDI19_2_/AtCALM1_2_ complex. **(F)** Sequence alignments of the ZZ domains from HsKCMF1 and AtUBR4. N-degron binding residues are highlighted with red circles. **(G)** MS analysis of the purified CeUBR4 complex with mitochondrial proteins highlighted in red and their intensity rank among all insect cell protein contaminants. **(H)** An AF3 model of ACADM monomer docked into the central EM density of the CeUBR4 complex.

We next examined species-specific adaptations of functional domains in the E4 complex. Notably, DI19 lacks an N-terminal ZZ domain, yet we co-purified the same substrate, SSBP1, as with the human complex and observed MTS-specific activity (**Fig. 5A,C**). These data can be explained by the presence of an internal ZZ domain in AtUBR4 (*49*), which shares high sequence similarity with the HsKCMF1 ZZ domain (**Fig. 5F**). Thus, while conserving the substrate targeting and ubiquitin conjugation mechanism, UBR4 complexes have reshuffled domains to generate distinct genetic contexts, facilitating the adoption of specialized PQC functions. Plants, for example, encode multiple homologues of the DI19 cofactor that respond to diverse stresses and localize to different cellular compartments (*55*). In *C. elegans*, the UBR4 complex appears to exhibit unique substrate selectivity, with mitochondrial medium-chain acyl-CoA dehydrogenase (ACADM) showing stronger enrichment compared to SSBP1 (**Fig. 5G, Table S4**), pointing to a distinct substrate filter. In line with this finding, we observed well-defined extra density in the central arena of CeUBR4, closely resembling the elongated shape of an orphaned subunit from the tetrameric ACADM complex (**Fig. 5H**). According to these cryo-EM data, bound ACADM not only contacts the ZZ domain but also two long helices in the N-terminal region. These two helices, which are unique to CeUBR4, protrude from the rigid E4 core into the central cavity, exposing a series of hydrophobic residues (**Fig. S7D**) that may help to recognize and bind the hydrophobic surface characteristic of orphaned subunits.

In conclusion, these findings reveal that despite evolutionary divergence, the E4 arena of UBR4 is remarkably well conserved across eukaryotes in its core architecture. However, evolutionary processes have reshuffled domains and interacting partners within the scaffold, producing holo E4 systems that are functionally similar but fine-tuned in their substrate profile and regulation for diverse PQC challenges in different organisms.

## Discussion

PQC enzymes target aberrant proteins for degradation based on broadly defined properties that are associated with protein dysfunction rather than on specific structural or sequence motifs. These properties often correlate with diverse phenotypes such as mislocalization, partial misfolding, orphan subunit status, or incomplete post-translational processing. Our structural, biochemical, and evolutionary analyses of UBR4 provide mechanistic insights into how such selection is achieved.

We demonstrate that the assembly of UBR4 with KCMF1 and CALM1 into an intricate 1.3 MDa E4 complex introduces important regulatory measures, ensuring tight control of degradation labelling beyond simple K48-ubiquitin chain extension. The giant structure of the E4 complex is composed of a rigid, ring-like core with flexibly attached functional domains. Substrates are sequestered within the central cavity, where they can interact with various receptor domains. The mobile E4 module protrudes to the narrow end of this arena, while the UBL domain is located between receptor domains and catalytic site, aligning ubiquitin moieties of captured substrates for chain extension. The bound co-factor KCMF1 has functional as well as structural roles in this assembly. Its C-terminal region stabilizes the core of the E4 complex, whereas the N-terminal ZZ-DZB motif protrudes into the center of the ring, acting as a substrate receptor. Moreover, the inherent mobility of the ZZ-DZB, located in the ubiquitin transfer region near the UBL domain, enables it to function as molecular bouncer, hindering ubiquitin chain extension of proteins that do not meet the specificity criteria. Accordingly, the ZZ-RING domain functions as a selectivity filter within the UBR4 system, identifying PQC substrates while preventing the erroneous marking of functional proteins for degradation.

UBR4 selects substrates based on their orphan protein character (*28*), the presence of an unprocessed MTS (*29*), or a specific N-degron (*56*). The weak sequence specificity of these selection criteria also explains the large number of reported UBR4 substrates. Efficient binding should depend on multivalent interactions, requiring multiple weak signals to be recognized. Additional contributions to substrate recognition likely come from peripheral domains like the BP, UBR and BS2, which also line the inner cavity. Moreover, for CeUBR4, we observed that two elongated helices in the N-terminus protrude into the ring and appear to bind the substrate, presumably recognizing the exposed hydrophobic surface present on an orphaned protein. Finally, the ring structure itself should enforce selectivity by restricting access to proteins below a certain size, favoring the binding of small orphaned proteins over multi-protein complexes. The functional importance of the ring architecture is emphasized by its conserved shape and dimension in CeUBR4, despite pronounced differences in its helical scaffold and dimerization interfaces. Moreover, structurally disruptive UBR4 patient mutations are spread throughout the ring scaffold suggesting the maintenance of this architecture is crucial for its PQC function (**Fig. S10**) (*57, 58*).

E4 ligases such as UBR4 play a dual role in amplifying ubiquitination signals and refining specificity in protein degradation (*59, 60*). Rather than relying solely on the priming E3 ligase for substrate selection, our findings emphasize the equally crucial role of E4 ligases in enhancing specificity. While both chain-initiating E3 ligases and chain-extending E4 ligases individually may exhibit weak substrate specificity, their sequential action yields an effective and specific proofreading mechanism to target aberrant proteins. Moreover, defective proteins are more efficiently targeted by E3/E4 ligase pairs the longer the substrates persist in a dysfunctional state. For example, mitochondrial precursor proteins imported efficiently into mitochondria escape recognition, but under stress conditions, stalled import traps them in the cytoplasm, where their chances of sequential recognition and degradation grow significantly. This temporal regulation should ensure the selective removal of truly defective proteins. Sequential ligase action also facilitates the formation of specialized PQC pathways towards distinct substrate types. For example, UBR4/KCMF1 operates alongside other general PQC enzymes, such as HUWE1, BIRC6, UBR5, HERC1 and HERC2 (*11, 30, 61–64*), each targeting distinct but overlapping sets of substrates. This combinatorial setting allows for fine-tuned regulation of PQC pathways in specific cell types and physiological conditions. Through its intricate structure, selective substrate targeting, and synergy with other PQC ligases, UBR4 is thus able to adopt a crucial role in cellular homeostasis, enabling the precise and efficient elimination of defective proteins.

## Acknowledgements

We would like to thank the EM facility of the Vienna BioCenter Core Facilities (VBCF), particularly H. Kotisch for collecting the cryo-EM datasets. We also thank the VBCF Protein Technologies facility and R. Imre from the VBCF Proteomics facility for their support. Finally, we thank all members of the Clausen lab for discussions.

This work was by the Austrian Research Promotion Agency Headquarter grant 852936 (TC), Marie Skłodowska-Curie grant 847548 (D.B.G.), Austrian Science Fund DocFunds grant DOC 112-B (J.F.E.), WWTF grants LS21-029 (L.E.B) and LS21-009 (L.D), and a BI fonds fellowship (L.E.B.). The IMP is supported by Boehringer Ingelheim. For the purpose of Open Access, the author has applied a CC BY public copyright licence to any Author Accepted Manuscript (AAM) version arising from this submission.

## Author Contributions

D.B.G and T.C. designed experiments. D.B.G., P.M., J.F.E., R.K., J.N. and L.D. performed cloning, expression and purification. D.B.G., P.M. and A.S. performed biochemical experiments. D.B.G. generated and analyzed structural data. A.S. performed the bioinformatic analysis. A.M., G.A.V., E.Z., R.S.H. and T.C. supervised research and provided scientific input. D.B.G. and T.C. coordinated the research project and prepared the manuscript with input from all authors.

## Materials and methods

### Cloning and expression

UBR4, KCMF1 and CALM genes from *H. sapiens*, *C. elegans* and *A. thaliana* were codon-optimized for insect cell expression and, where required, synthesized in fragments flanked with BsaI sites. UBR4 was synthesized with a C-terminal His_10_ tag, KCMF1 with a C-terminal StrepII tag and CALM with an N-terminal FLAG tag. The three genes were assembled via scarless GoldenGate reactions into a pGBdest vector with each gene flanked by a polyhedrin promoter and an SV40 polyA signal (*65*). For monomeric HsUBR4 and CeUBR4 the same procedure was used except the His_10_ tag was replaced by a StrepII tag. Mutations were generated by Gibson assembly. The plasmid was amplified in three equally sized fragments with complementary overhangs. Mutations were introduced in one fragment using a blunt-end ligation strategy. The sequence of all generated insect cell constructs was confirmed by whole plasmid sequencing. Plasmids were transformed into DH10EmBacY cells and blue-white screening was used to select colonies with bacmids. Bacmids were extracted by alkaline lysis and isopropanol precipitation and then transfected using Polyethylenimine PEI 25K into Sf9 cells (Expression Systems) for viral amplification. Protein expression was performed in Trichoplusia ni High-Five cells (Thermo Fisher). Cells were infected with virus at a density of 1.5 x 10^6^ ml^-1^ and grown for three days at 27°C. Cells were harvested by centrifugation at 600 x g.

Constructs for E. coli expression were generated by Gibson assembly. His_6_-SUMO-MTS1/2-linker-UbΔGG was synthesized with overhangs and inserted into a pET28 vector. The linker comprised 21 glycine or serine residues. The control construct with no MTS as well as N-degron versions were generated by blunt-end mutagenesis targeting the region directly after the SUMO cleavage site and before the linker. Plasmids were transformed into BL21 (DE3) cells for expression in Lysogeny Broth media. Expression was performed at 17°C with 0.2 mM isopropyl-ß-d-thiogalactopyranoside.

### Protein purification

HsUBR4/KCMF1/CALM1 and AtUBR4 expression pellets were resuspended in buffer containing 50 mM HEPES pH 7.5, 500 mM NaCl, 0.5 mM TCEP, 20 mM imidazole with benzonase (MBS) and a Complete EDTA-free protease inhibitor tablet (Roche). Cells were lysed using a glass douncer and cleared by centrifugation at 40,000 x g. The soluble lysate was loaded on a 5 mL HisTrap HP (Cytiva) column using an Akta PURE system (Cytiva). The column was washed with 7 column volumes (CVs) of the lysis buffer followed by 6 CVs of buffer with 80 mM imidazole. UBR4 complexes were then eluted with a linear gradient from 80 to 300 mM imidazole. SDS-PAGE was used to identify fractions containing UBR4. For the HsUBR4/KCMF1/CALM1 co-expression construct, the eluted protein was concentrated by ultrafiltration to 2 mL and subjected to size-exclusion chromatography using a Superose 6 pg 16/70 column (Cytiva) equilibrated in 20 mM HEPES pH 7.5, 250 mM, 0.5 mM TCEP. Fractions containing HsUBR4/KCMF1/CALM1 were concentrated by ultrafiltration, flash frozen and stored at −70°C. AtUBR4 was purified by anion-exchange chromatography using a 6 mL Resource Q column (Cytiva) instead of SEC after the His-affinity purification in 50 mM HEPES pH 7.5, 0.5 mM TCEP and a 150-1000 mM NaCl gradient. Fractions containing AtUBR4 were pooled, concentrated and flash frozen.

For the CeUBR4/KCMF1 complex, lysis followed the same protocol except imidazole was excluded from the buffer. The clarified lysate was first purified by Strep-affinity chromatography using a 5 mL StrepTrap HP column (Cytiva) equilibrated in lysis buffer. The column was washed with 8 CVs of the same buffer and then eluted in the same buffer with 2.5 mM desthiobiotin. The complex was subjected to SEC using the same protocol as for HsUBR4/KCMF1/CALM1. Following SEC, the complex was loaded on a 1 mL HisTrap HP column, washed with 20 mM and then 50 mM imidazole and eluted with 300 mM imidazole. The imidazole was then removed by repeated concentration and dilution using a spin concentrator and flash frozen and stored at −70°C.

For HsUBR4 and CeUBR4 monomeric complexes, cell pellets were resuspended in phosphate buffered saline with 0.5 mM TCEP (PBS-TCEP) and lysed and subjected to Strep-affinity chromatography as above. The proteins were further purified by anion-exchange chromatography using a 6 mL Resource Q column (Cytiva) in PBS-TCEP with and a 250-500 mM NaCl gradient. Fractions containing UBR4 were pooled, concentrated and flash frozen.

Expression pellets for the SUMO-linker-Ub* constructs were resuspended in 50 mM Tris-HCl pH 8.0, 500 mM NaCl, 0.5 mM TCEP, 25 mM imidazole, benzonase and protease inhibitors and lysed by sonication. Clarified lysate was applied to a 5 mL HisTrap HP (Cytiva) using a syringe, washed with 5 CVs of the same buffer and then eluted with 250 mM imidazole in the same buffer. The eluate was treated with his_6_-SENP2 protease (MBS) overnight at 4oC to remove the SUMO and expose the N-degron. The imidazole concentration was reduced to 25 mM imidazole by dilution in 50 mM Tris-HCl pH 8.0, 500 mM NaCl, 0.5 mM TCEP and the sample was reapplied through the 5 mL HisTrap HP column using a syringe to remove his_6_-SUMO and his_6_-SENP2. The column was washed with an additional 3 CVs of buffer with 25 mM imidazole and all flow-through collected, concentrated and then flash frozen and stored at −70°C.

UBE2D2, UBE2A, UBA1, and ubiquitin were purified as previously described (*8, 63*). The concentration of all proteins was determined by absorbance at 280 nm using calculated extinction coefficients.

### Cryo-EM grid preparation and data collection

1.3 mg/ml HsUBR4/KCMF1/CALM1 or CeUBR4/KCMF1 in 25 mM HEPES pH 7.5, 200 mM NaCl, 0.5 mM TCEP was applied onto a freshly glow-discharged (90 seconds at 25 mA) Quantifoil R1.2/1.3 Cu 200 mesh grid. Grids were blotted for 1.2 seconds before rapid freezing in liquid ethane using a Leica GP2 plunge-freezer. For the HsUBR4/KCMF1/CALM1/UBE2A complex the protein was first incubated with 3 μM UBE2A (a 1.5 fold excess). For EGTA treatment the sample was first incubated with 2 mM EGTA for 30 minutes. For all cases except one HsUBR4/KCMF1/CALM1 dataset, 0.8 mM CHAPSO was added to the sample directly before freezing. A Glacios TEM (Thermo Fisher) equipped with a Falcon 3 detector was used to screen grids. For data collection, the grids were transferred to a Titan Krios G4 (Thermo Fisher) with a Falcon 4EC detector operated by the Research Institute of Molecular Pathology, Austria. All data were collected with the same parameters using the EPU software (Thermo Fisher) with 130,000x magnification (0.951 Å pixel size) and a total dose of 50 e/Å^2^. Patch Motion correction was performed on-the-fly using CryoSPARC Live (*66*).

### Cryo-EM data analysis

The HsUBR4/KCMF1/CALM1 structure was generated by combining three different datasets – one without CHAPSO, one with CHAPSO and one with CHAPSO and EGTA. The HsUBR4/KCMF1/CALM1/UBE2A and CeUBR/KCMF1 structures were from single datasets. All datasets were processed using the same strategy with specifics shown in Figs S2, S3 and S5. Motion corrected micrographs were simultaneously imported into CryoSPARC v4 (*66*) and Relion 4.0 (*67*). CTF correction, particle picking and removal of junk particles was performed independently with both programs. In CryoSPARC, patch CTF estimation followed by blob picking was used to generate templates for template picking. After template picking, particles were extracted in a 512 pixel box size downscaled to 128 pixels. Many rounds of 2D classification were performed until only particles with clear secondary structure remained. All initial models were generated by Ab Initio in CryoSPARC. In Relion 4, after CTF estimation with CTFFIND4 (*68*), 1000 particles were manually picked to generate templates for autopicking. Autopicked particles were extracted at 512 pixels and downscaled to 100 pixels. Repeated round of 2D classification were used to only remove ice and edges. 3D classification with alignment with the CryoSPARC initial model was then used to remove low quality particles.

These two particle sets were then combined in CryoSPARC, duplicates were removed and then extracted with a box size of 512 pixels, downscaled to 384 pixels (1.27 Å/px). The extracted particle stack was then imported into Relion for all further processing. To obtain clear density for the co-purified substrate in the CeUBR4/KCMF1 structure a 3D classification with alignment was performed. For all other maps, first a 3D refinement was performed with either C1 or C2 symmetry enforced. The particles refined with C2 symmetry were subjected to symmetry expansion. Then soft masks were generated for the various parts of the structure and 3D classification without alignment was used using either the C1, C2 or symmetry expanded aligned particles and different T values to find the best particle sets for each region of the structure. These particles were subjected to either global or focused refinement to obtain the final maps. A composite map was made in ChimeraX (*69*) by aligning the focused maps onto a single map which had all features moderately well resolved. Certain regions were selected based on the molecular model, scaled and combined with the volume maximum command.

### Molecular model building and refinement

The focused maps were used for all initial modelling, with these models then combined into the composite. The C-terminal and N-terminal dimerization regions of the HsUBR4/KCMF1/CALM1 structure were autobuilt using ModelAngelo (*70*). For the CeUBR4/KCMF1 structure, ModelAngelo was used to build the C-terminal dimerization region as well as the region comprising the BP domain and associated ARM repeats. Additionally, many AlphaFold3 models were predicted for all sections of the structure including the various protein-protein and protein-zinc interactions using the AlphaFold3 server (*37*). These were generally accurate enough at a local level to be docked directly and confidently in the map. However, they differed significantly from the real structure at a global conformational level. From these starting models we used rounds of modelling in Coot (*71*) and real space refinement in Phenix Real Space Refine (*72*) to obtain models which fit well to the density while having reasonable geometry statistics. Model to map fits are shown in Fig. S3 and S9 while refinement statistics are shown in Table S1.

### Negative stain EM

Continuous carbon coated copper/palladium grids (Agar Scientific) were glow discharged at 25 mA for 60 seconds using a BAL-TEC sputter coater CD005. AtUBR4 was diluted to 0.05 mg/mL in 25 mM HEPES pH 7.5, spotted on the grids, incubated for one minute, blotted, washed with 4 μL 2% uranyl acetate, blotted again, and incubated for one minute with 4 μL 2% uranyl acetate before final blotting. Data were cp;;ected on a 200 kV FEI Tecnai G2 20 (Thermo Fisher Scientific) at 50,000x magnification with a 2.21 Å pixel size using SerialEM. Particle picking was performed using CrYOLO and then particles coordinates were imported into CryoSPARC v4 for extraction with a 2x binned 256 pixel box size and then rounds of 2D classification.

### Ubiquitination assays

All ubiquitination assays were performed in buffer containing 25 mM HEPES pH 7.5, 150 mM NaCl, 0.5 mM TCEP, 5 mM MgCl_2_. Reactions contained 100 or 200 nM UBR4 monomer as indicated, 250 nM UBA1, 500 nM UBE2A, 5 μM substrate-Ub* and either 5 μM Ub-K0 or 10 uM wild-type ubiquitin spiked with 1 μM Ub-DyLight488. Reactions were initiated by the addition of 5 mM ATP and proceeded for 15, 30 or 60 minutes as indicated.

Reactions were quenched with SDS gel loading buffer and separated by SDS-PAGE using 4-12% NuPAGE Bis-Tris gradient gels (Invitrogen) in MES running buffer. If required, gels were first imaged for DyLight488 fluorescence using a ChemiDoc MP system (Bio-Rad) before Coomassie staining. All experiments were replicated entirely at least twice.

### Mass photometry

Experiments were performed using a OneMP mass photometer (Refeyn Ltd.) controlled by the AcquireMP application. A one-minute acquisition time in the medium field of view was used. To obtain average species masses, particle event histograms were automatically Gaussian fitted with the DiscoverMP software. Samples were first prepared in a 200 nM stock solution in 25 mM HEPES pH 7.5, 150 mM NaCl, 0.5 mM TCEP and prior to measurement were diluted 10x in PBS. Data were plotted in the DiscoverMP application.

### Co-immunoprecipitation and mass spectrometry experiments

Co-IP/MS experiments were performed using the lysate of RKO-Cas9 cells with a doxycycline-inducible *UBR4*-targeting sgRNA as previously described in detail (*62, 63*). UBR4 KO was confirmed by Western blot. Cells were washed and resuspended in PBS before lysis by sonication. In addition to PBS the lysis buffer contained proteasome inhibitor (100 μM MG-132) 1 mM PMSF, benzonase and 0.1% NP40. After clarification by centrifugation the total protein concentration was determined using the BCA assay. All lysates were normalized to 1 mg/ml total protein. For lysates which were not treated with deubiquitinase, the deubiquitinase inhibitor PR619 was added at 100 μM. Experiments were performed in triplicate. For deubiquitinase treatment 250 nM USP21 was used and deubiquitination confirmed by Western blot. MagStrep Strep-Tactin beads (IBA) were pre-bound to either UBR4 or GFP by incubation at 4°C for one hour followed by washing with PBS. 7 μL of bead slurry was added to 550 μL of treated RKO lysate and incubated overnight at 4°C. Beads were washed twice with 500 μL PBS before resupension in 100 μL PBS for mass spectrometry analysis.

Samples were treated with trypsin in ammonium bicarbonate before extraction with 5% formic acid. Peptide desalting was performed using an Oasis HLB 96-well µElution Plate with 2 mg Sorbent (Waters) and organic content of the eluates were removed by evaporation in a vacuum centrifuge.

The nano HPLC system used was an UltiMate 3000 RSLC nano system (Thermo Fisher Scientific) coupled to a Q Exactive HF-X mass spectrometer (Thermo Fisher Scientific), equipped with a Proxeon nanospray source (Thermo Fisher Scientific). Peptides were loaded onto a trap column (Thermo Fisher Scientific, PepMap C18, 5 mm × 300 μm ID, 5 μm particles, 100 Å pore size) at a flow rate of 25 μL min^-1^ using 0.1% TFA as mobile phase. The trap column was switched in line with the analytical column (Thermo Fisher Scientific, PepMap C18, 500 mm × 75 μm ID, 2 μm, 100 Å) after 10 minutes and peptides were then eluted using a binary one hour gradient, starting from 98% A (water/formic acid, 99.9/0.1, v/v) and 2% B (water/acetonitrile/formic acid, 19.92/80/0.08, v/v/v), increasing to 35% B over 60 minutes and then to 95% B over 5 minutes with a flow rate of 230 nl min^-1^.

The Orbitrap Exploris 480 mass spectrometer was operated in data-dependent mode, performing a full scan (m/z range 350-1200, resolution 60,000, normalized AGC target 100%) at 3 different compensation voltages (CV −45, −60, −75), followed each by MS/MS scans of the most abundant ions for a cycle time of 1 second per CV. MS/MS spectra were acquired using HCD collision energy of 30, isolation width of 1.2 m/z, orbitrap resolution of 30,000, normalized AGC target 200%, minimum intensity of 50,000 and maximum injection time of 100 ms. Precursor ions selected for fragmentation (include charge state 2-6) were excluded for 20 s. The monoisotopic precursor selection (MIPS) filter and exclude isotopes feature were enabled.

For peptide identification, the RAW-files were first loaded into Proteome Discoverer (version 2.5.0.400, Thermo Scientific). MS/MS spectra were then searched using MSAmanda v2.0.0.19924 (*73*). The fragment and peptide mass tolerance were set to ±10 ppm and the maximum number of missed cleavages was set to 2, using tryptic enzymatic specificity without proline restriction. Peptide and protein identification was initially performed by searching the RAW-files against the Uniprot-database using the taxonomy Trichoplusia ni (2021-03; 21,163 sequences; 13,598,141 residues) and Homo sapiens (2023-03; 20,516; 11,401,144 residues) supplemented with common contaminants and the sequences of UBR4, KCMF1 and CALM1 using Beta-methylthiolation on cysteine as a fixed modification. The result was filtered to 1 % FDR on protein level using the Percolator algorithm (*74*) integrated in Proteome Discoverer. Following this, a second round of searching was done on the sub-databases of these proteins looking for protein modifications and again filtered. Peptides were subjected to label-free quantification using IMP-apQuant (*75*). Proteins were filtered to be identified by a minimum of 2 PSMs in at least 1 sample. Identified proteins were pre-filtered to contain at least 3 quantified peptide groups.

## Data availability

Structure coordinates have been deposited in the Protein Data Bank (PDB) under the accession codes listed in **Table S1**. Cryo-EM maps have been deposited in the Electron Microscopy Data Bank (EMDB) under accession codes listed in **Table S1**. All other data are provided with this paper or available upon request.

## Supplementary Figures legends

**Fig. S1.**
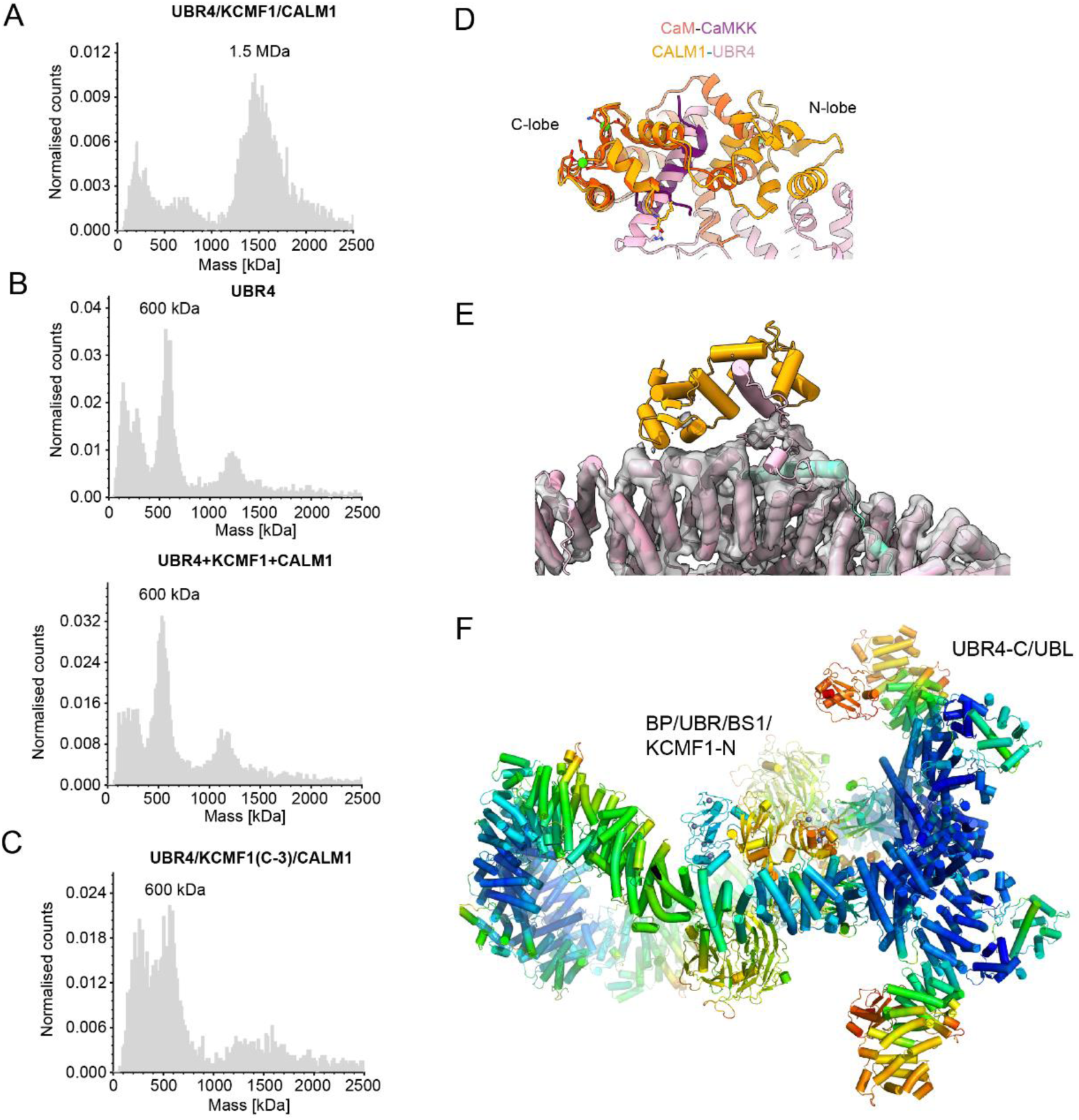
Assembly and structure of the HsUBR4/KCMF1/CALM1 complex. **(A-C)** Mass photometry analysis of **(A)** the co-expressed complex, **(B)** UBR4 alone and UBR4 mixed with isolated KCMF1 and CALM, and **(C)** the UBR4/KCMF1-3C/CALM1complex. **(D)** Comparison of the HsUBR4-HsCALM1 interaction with a canonical CALM interaction (calmodulin-calmodulin dependent protein kinase kinase, PDB 1IQ5). **(D)** Mass photometry analysis of HsUBR4(ΔCIH)/KCMF1/CALM1. **(E)** Model of the HsUBR4/KCMF1/CALM1 complex docked into a map where one copy of CALM1 is absent. **(F)** Cryo-EM model of the HsUBR4/KCMF1/CALM1 complex colored by B factor to indicate the flexibility of different regions. Red indicates a higher B factor.

**Fig. S2.**
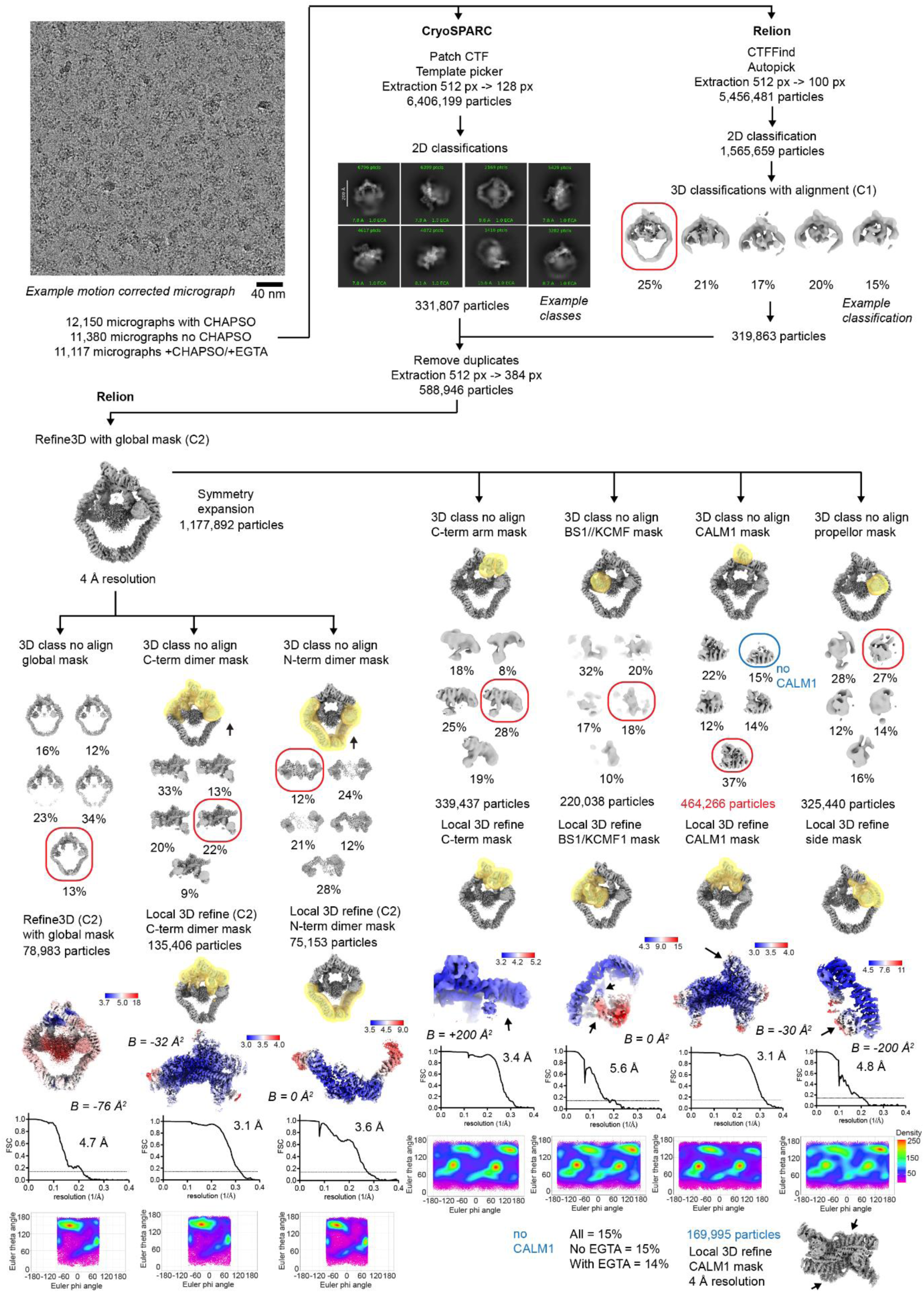
Cryo-EM processing pipeline for the HsUBR4/KCMF1/CALM1 complex. Further investigation of CALM1 binding is indicated.

**Fig. S3.**
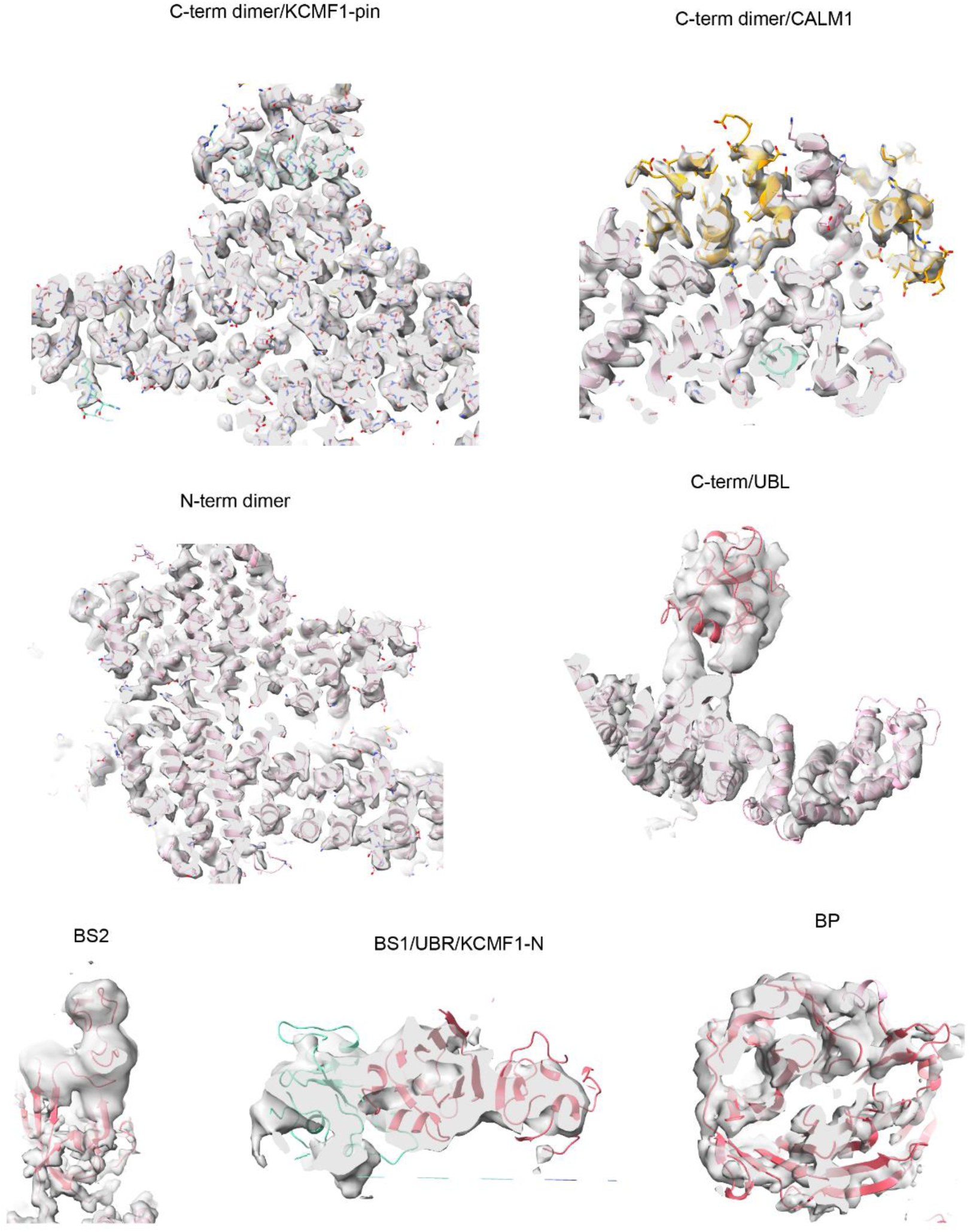
Example map to model fits for different regions of the HsUBR4/KCMF1/CALM1 structure.

**Fig. S4.**
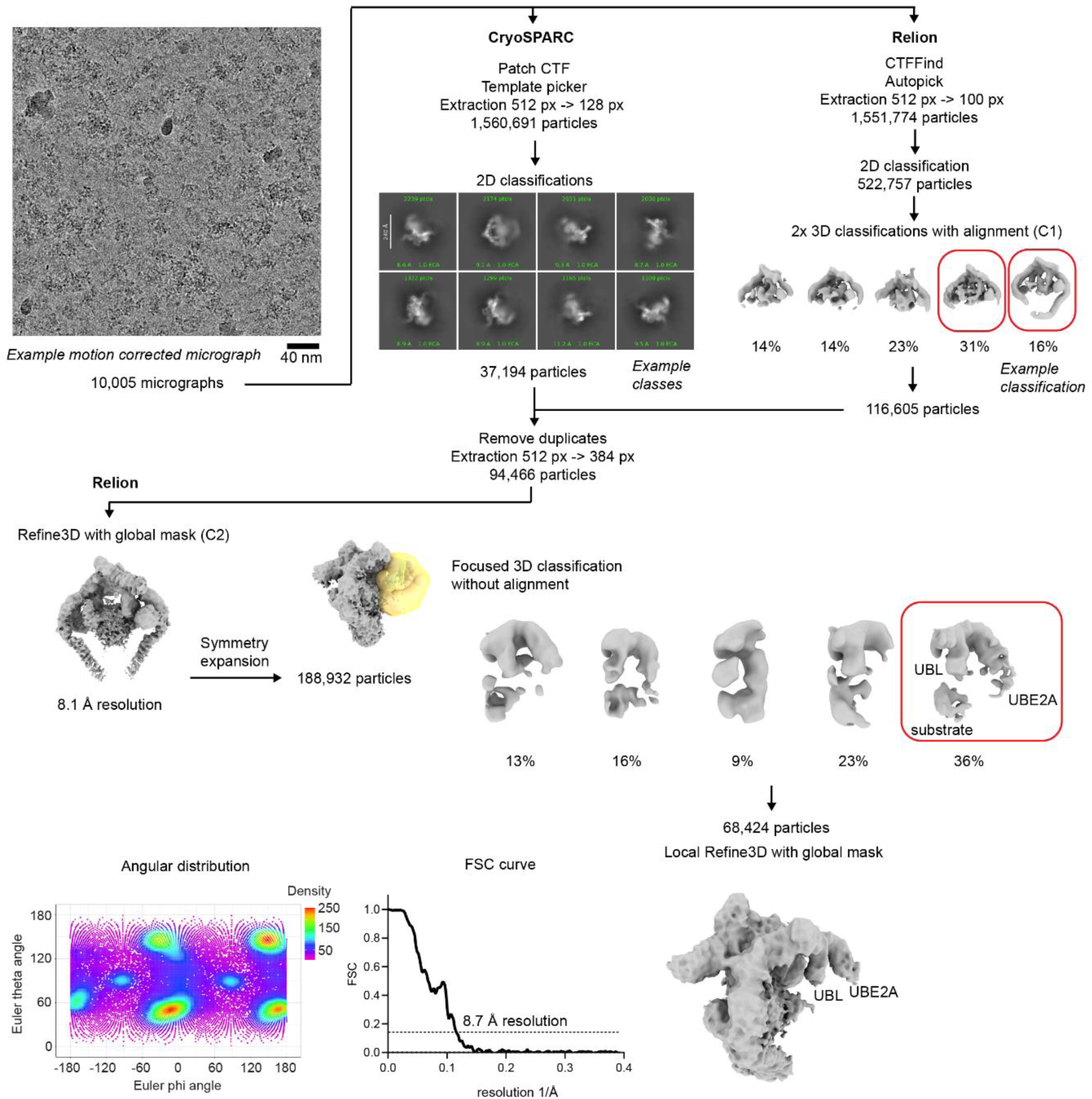
Cryo-EM processing pipeline for the HsUBR4/KCMF1/CALM1 complex incubated with HsUBE2A.

**Fig. S5.**
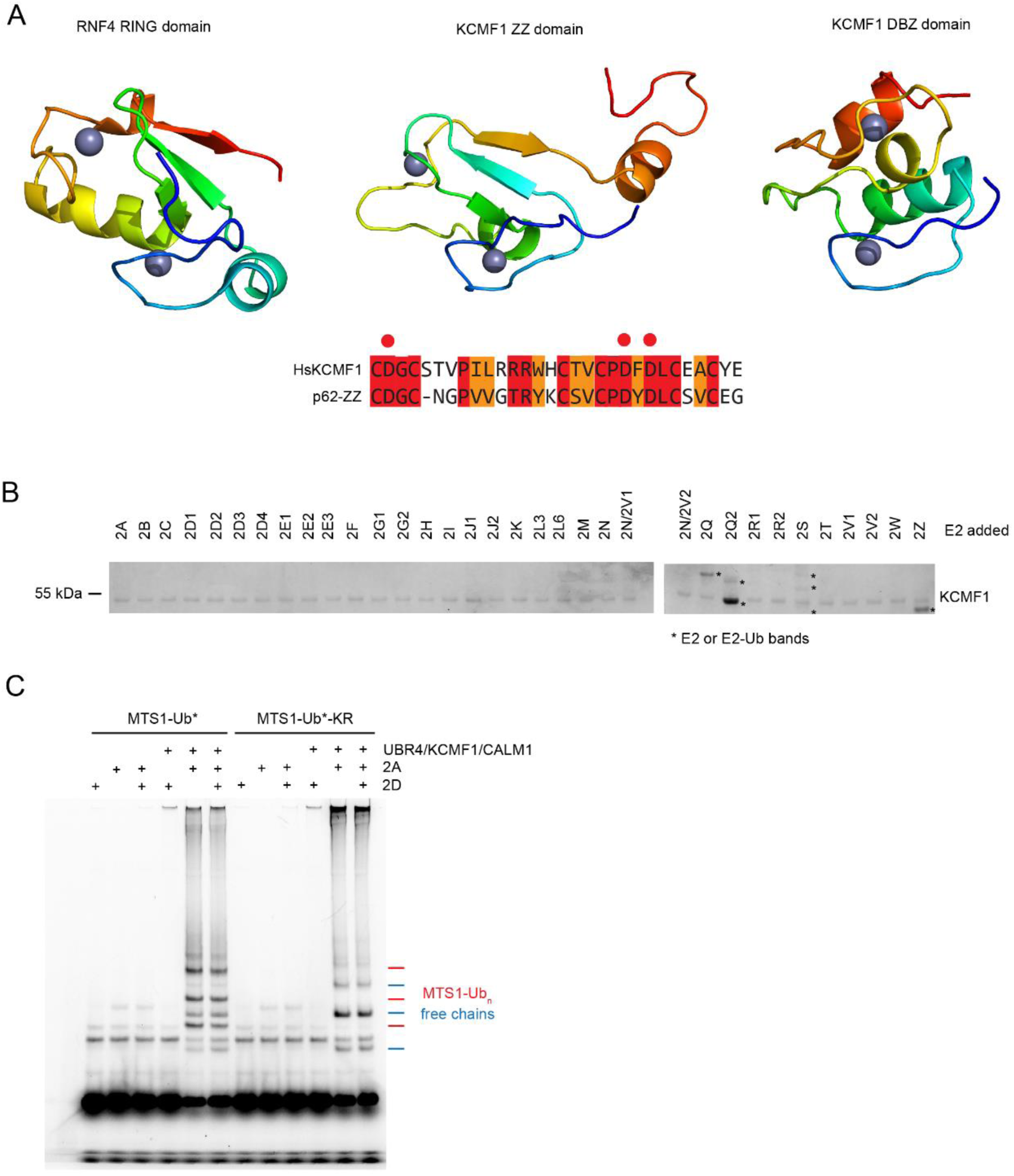
KCMF1 is not an E3 ligase. **(A)** Structural and sequence analysis of the N-terminal Zn-binding domains of KCMF1 with the RING domain of RNF4 (PDB 4AP4) as a reference. ZZ domain N-degron binding residues are highlighted in the alignment. **(B)** E2 scan kit assay. HsKCMF1 was incubated with the indicated E2 enzymes along with Ub, UBA1 and ATP for 60 minutes at 37°C. Indicated bands are also present in the absence of HsKCMF1. **(C)** Ubiquitination assay with 200 nM HsUBR4/KCMF1/CALM1 and the indicated Ub* substrates. Here, wild-type dylight-488 ubiquitin was used as a substrate generating either free chains or chains on the Ub* substrate. Reactions included either UBE2A, UBE2D2 or both E2 enzymes.

**Fig. S6.**
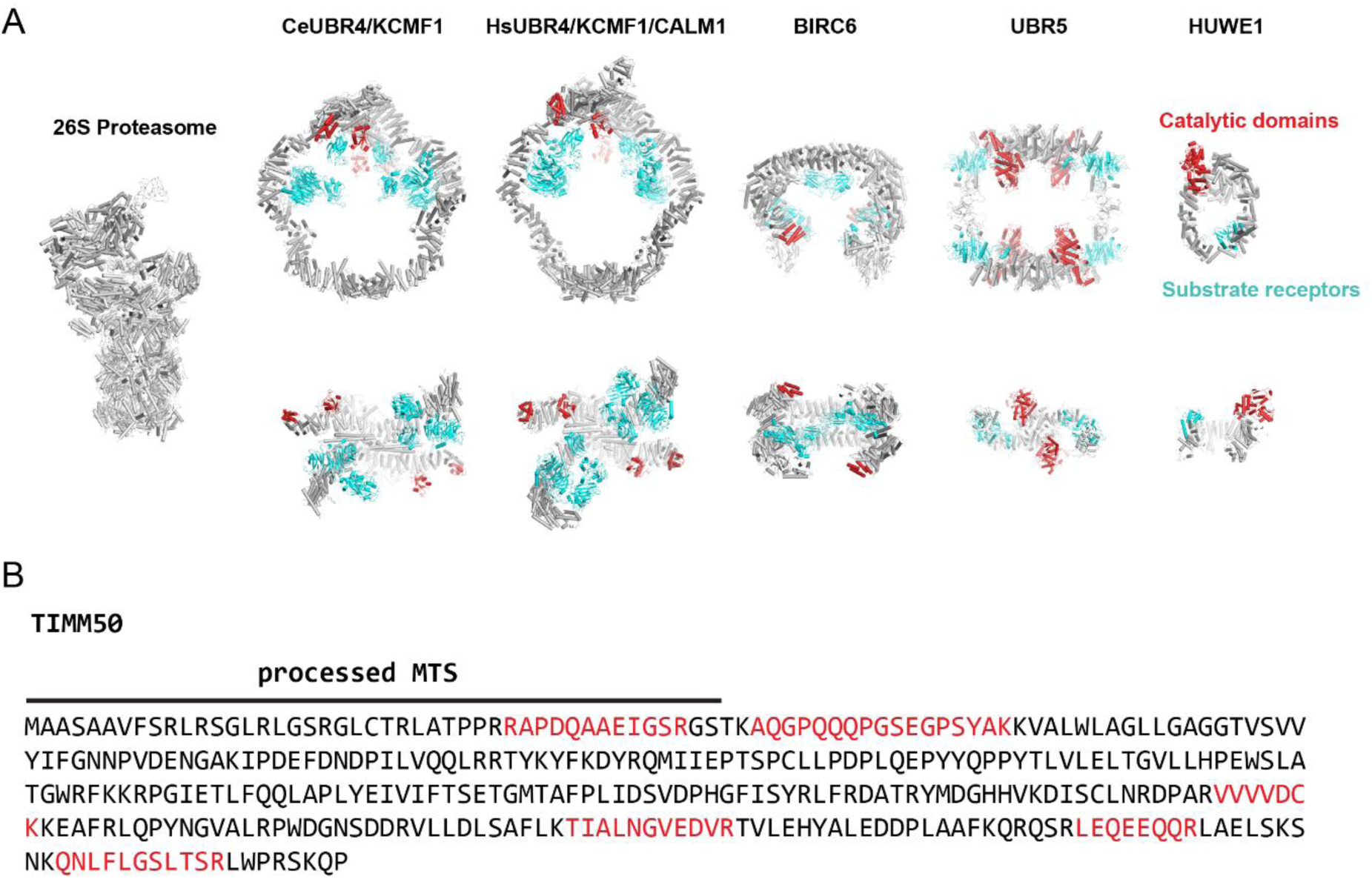
Substrate recognition by the UBR4 complex. **(A)** Comparison of the human UBR4 complex with other giant E3 ligases. The 26S proteasome is used as a size reference. Catalytic domains and putative substrate receptor domains are indicated in red and cyan respectively. Models used are: HUWE1 (PDB 7JQ9), UBR5 (PDB 8EWI), BIRC6 (PDB 8ATU) and the proteasome (PDB XXX). **(B)** Example protein found in the UBR4 co-IP/MS analysis with confidently identified peptides shown in red and the position of the processed MTS indicated.

**Fig. S7.**
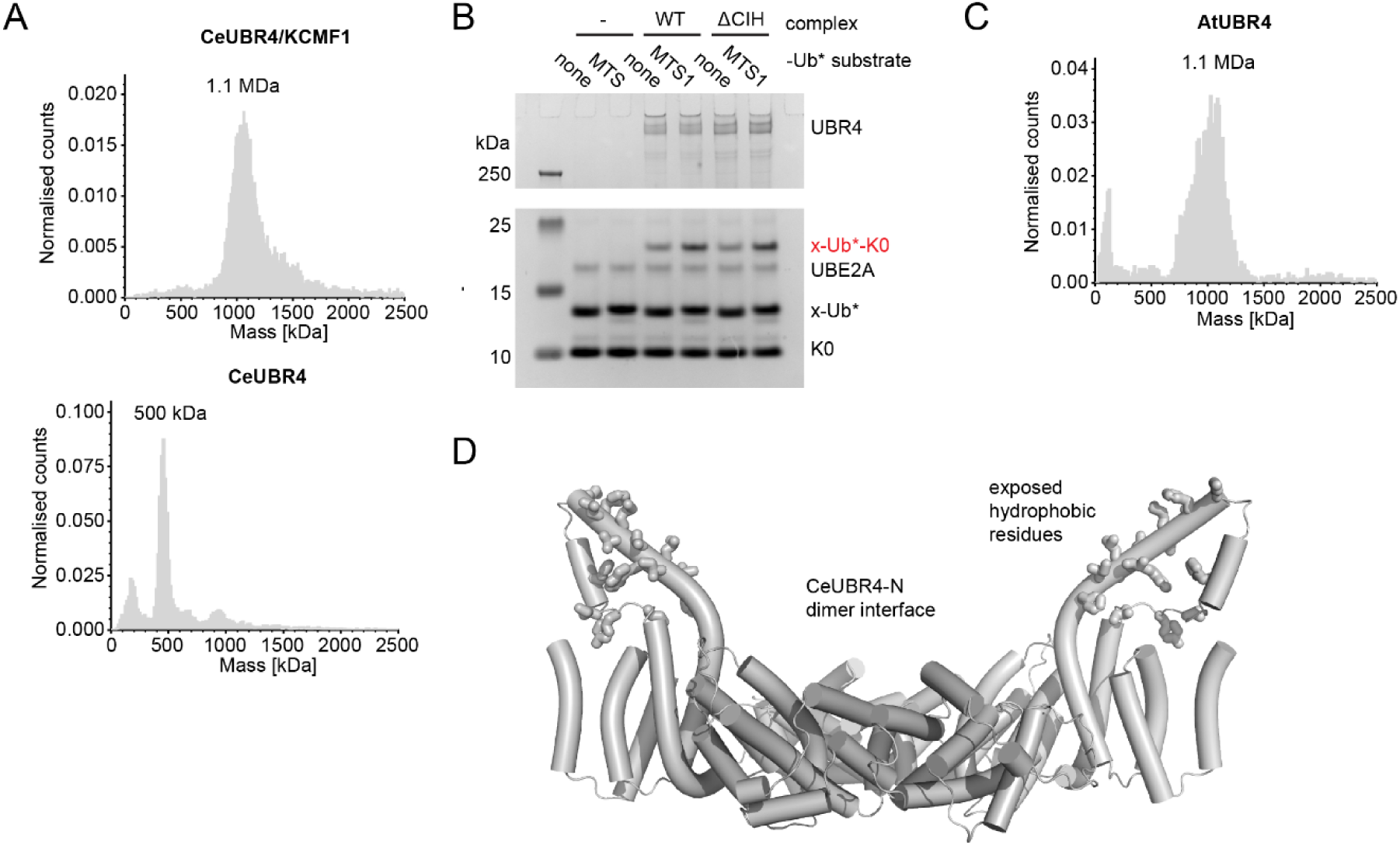
Evolutionary analysis of the UBR4 complex. **(A)** Mass photometry analysis of the CeUBR4 complex and CeUBR4 alone. **(B)** E4 ligase assay with the indicated human UBR4 complex (100 nM) and Ub* substrate for 45 minutes at 37°C. **(C)** Mass photometry analysis of AtUBR4. **(D)** AlphaFold3 model of the CeUBR4 N-terminal dimer interface. All the exposed hydrophobic residues on the two helices which enter into the central arena are shown in stick representation. These helices were not fully resolved in the cryo-EM structure due to flexibility.

**Fig. S8.**
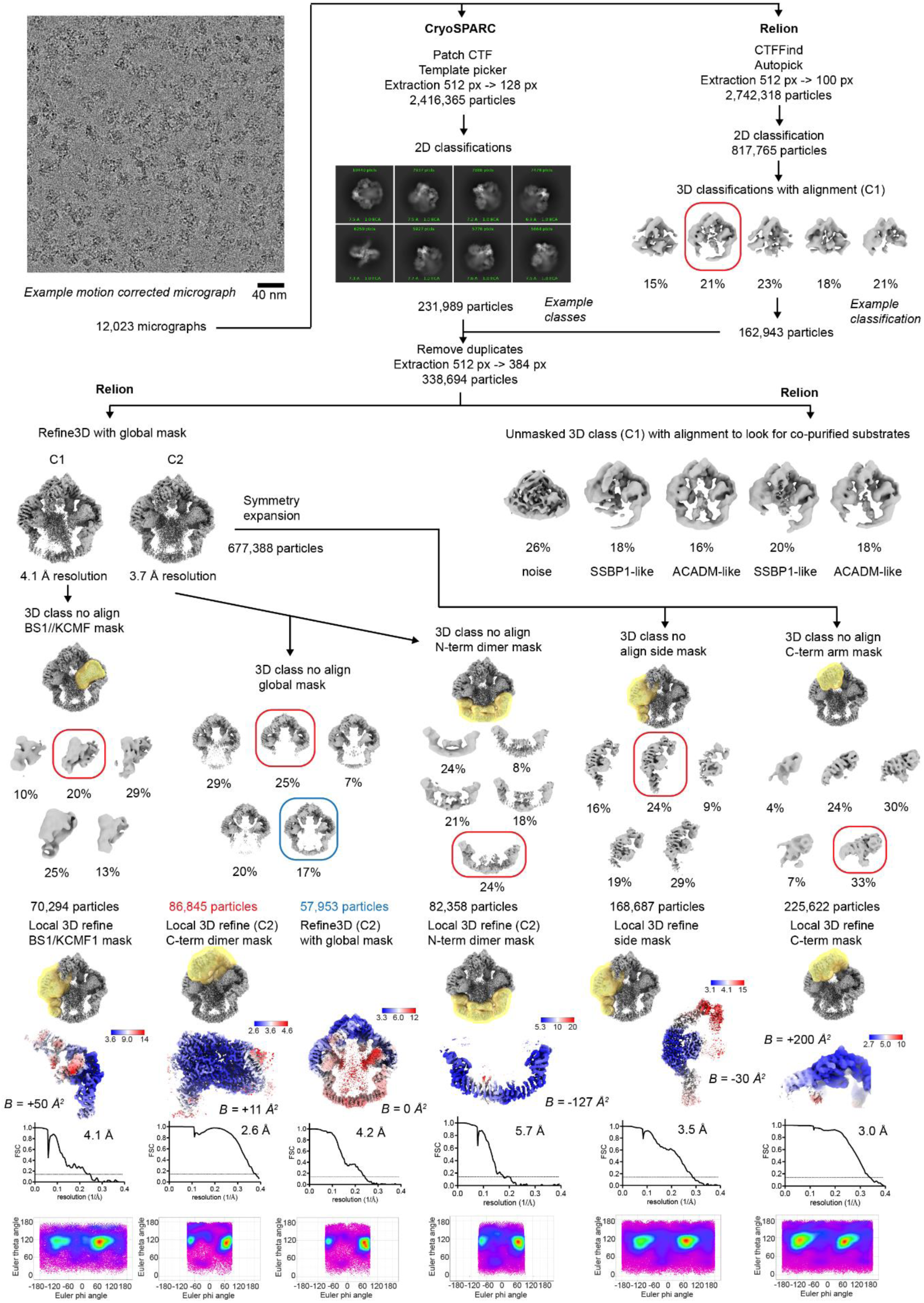
Cryo-EM processing pipeline for the CeUBR4 complex. Analysis of the different substrate bound states is shown in the center-right of the pipeline.

**Fig. S9.**
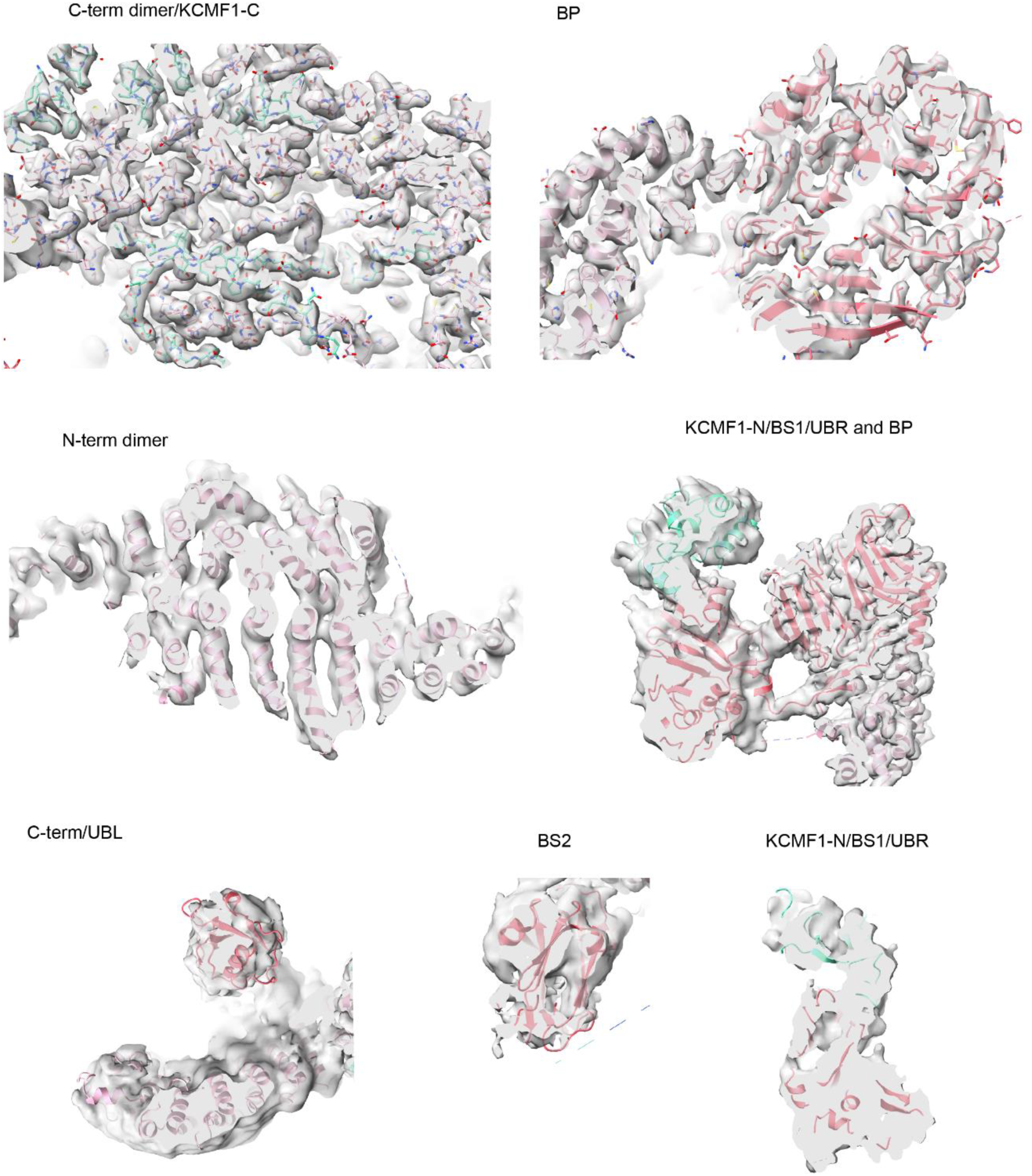
Quality of the density and model for the CeUBR4 complex in different regions of the structure.

**Fig. S10.**
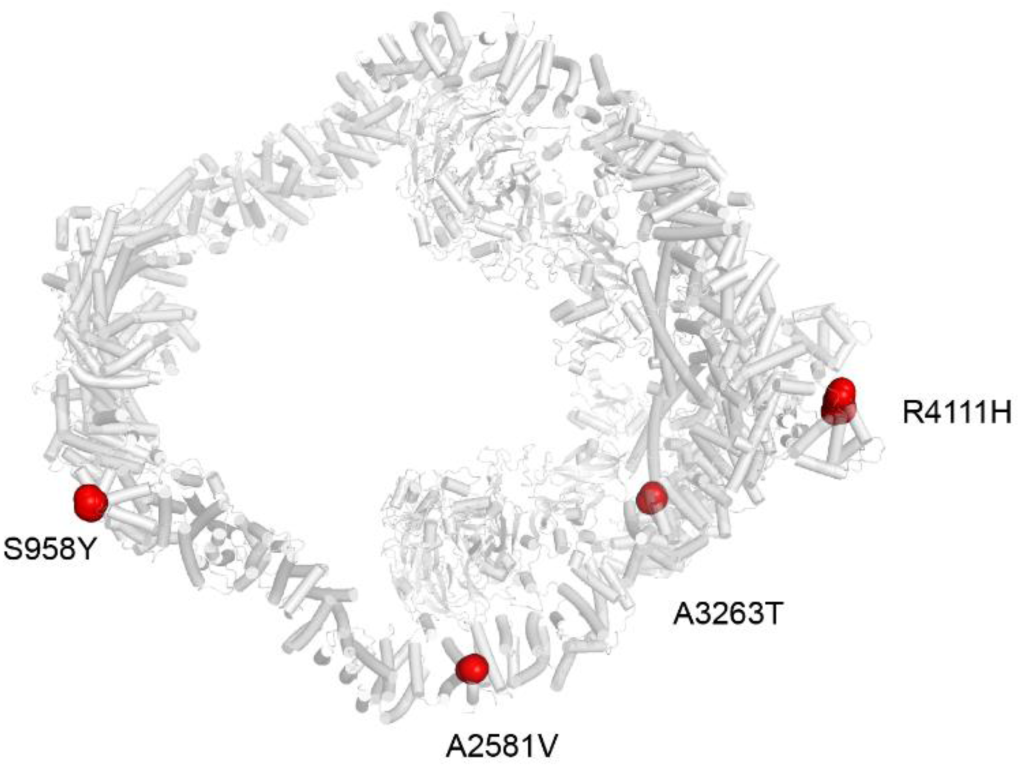
UBR4 patient mutations within the Armadillo repeats. Mutated residues are shown as red spheres.

**Fig. S11.**
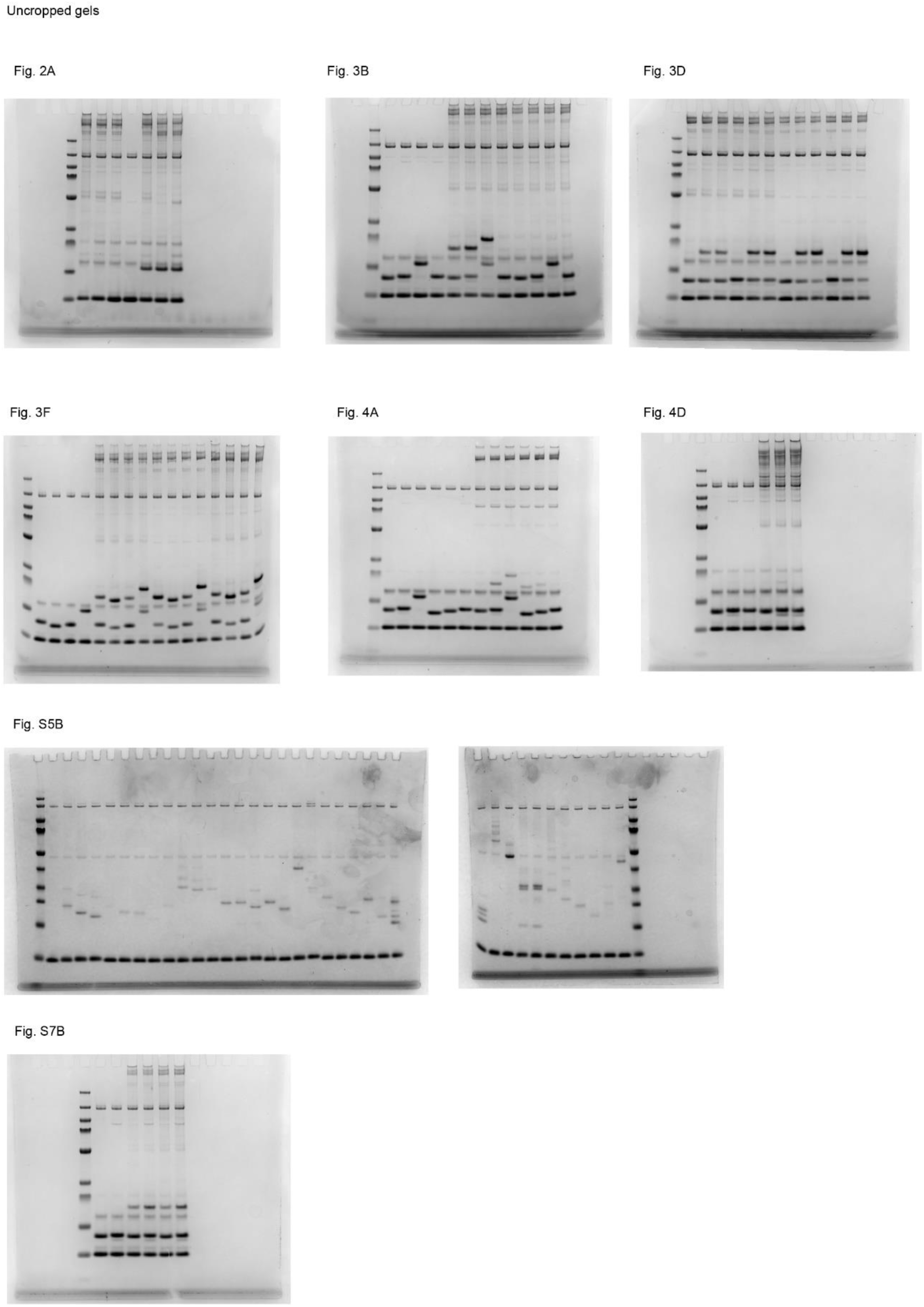
Uncropped gels.

## Supplementary Tables

**Table S1.** Cryo-EM data collection and refinement statistics. See Fig. S2 for details. *Resolution range includes unmodelled substrate density.

|  |  |  |  |
| --- | --- | --- | --- |
|  | HsUBR4/KCMF1<br>CALM1<br>PDB XXXX, EMD-XXXXX<br>EMD-XXXXX<br>EMD-XXXXX<br>EMD-XXXXX<br>EMD-XXXXX<br>EMD-XXXXX<br>EMD-XXXXX | HsUBR4/KCMF1/<br>CALM1/UBE2A<br>EMD-XXXXX | CeUBR4/KCMF1<br>PDB XXXX, EMD-XXXXX<br>EMD-XXXXX<br>EMD-XXXXX<br>EMD-XXXXX<br>EMD-XXXXX<br>EMD-XXXXX |
| Magnification | 130,000x | 130,000x | 130,000x |
| Voltage (kV) | 300 | 300 | 300 |
| Electron exposure (e<sup>-</sup>/Å<sup>2</sup>) | 50 | 50 | 50 |
| Defocus range (μm) | -0.8 – -2.0 | -0.8 – -2.0 | -0.8 – -2.0 |
| Pixel size (Å) | 0.951->1.27 | 0.951->1.27 | 0.951->1.27 |
| Symmetry imposed | C2 | C1 | C2 |
| Initial particle images (no.) | 6,406,199+<br>5,456,481 | 1,560,691+<br>1,551,774 | 2,416,365+<br>2,742,318 |
| Final particle images (no.) | 78,983+<br>135,406+<br>75,153+<br>339,437+<br>220,038+<br>464,266+<br>325,440 | 68,424 | 70,294+<br>86,845+<br>57,953+<br>82,358+<br>168,687+<br>225,622 |
| Map resolution (Å) | 4.7, 3.1, 3.6, 3.4,<br>5.6, 3.1, 4.8 | 8.7 | 4.1, 2.6, 4.2, 5.7,<br>3.5, 3.0 |
| FSC threshold | 0.143 | 0.143 | 0.143 |
| Map resolution range* | 3.0-30 | 6.5-30 | 2.6-30 |
| Initial model used | AlphaFold3 |  | AlphaFold3 |
| Model resolution (Å) | 3.3 |  | 3.1 |
| FSC threshold | 0.5 |  | 0.5 |
| Map sharpening B factor (Å<sup>2</sup>) | See Fig. S2 |  | See Fig. S8 |
| Non-hydrogen atoms | 68,966 |  | 64,704 |
| Protein residues | 8,794 |  | 8058 |
| Ligands | 20 x Zn<sup>2+</sup> |  | 18 x Zn<sup>2+</sup> |
| *B* factors (Å<sup>2</sup>) |  |  |  |
| Protein | 86 |  | 107 |
| Ligand | 111 |  | 116 |
| r.m.s.d. |  |  |  |
| Bond lengths (Å) | 0.010 |  | 0.006 |
| Bond angles (°) | 0.835 |  | 0.767 |
| MolProbity score | 1.99 |  | 1.98 |
| Clashscore | 14.7 |  | 12.0 |
| Poor rotamers (%) | 1.4 |  | 1.5 |
| Ramachandran |  |  |  |
| Favored (%) | 96.70 |  | 96.28 |
| Allowed (%) | 3.23 |  | 3.66 |
| Disallowed (%) | 0.07 |  | 0.06 |

**Table S2.**
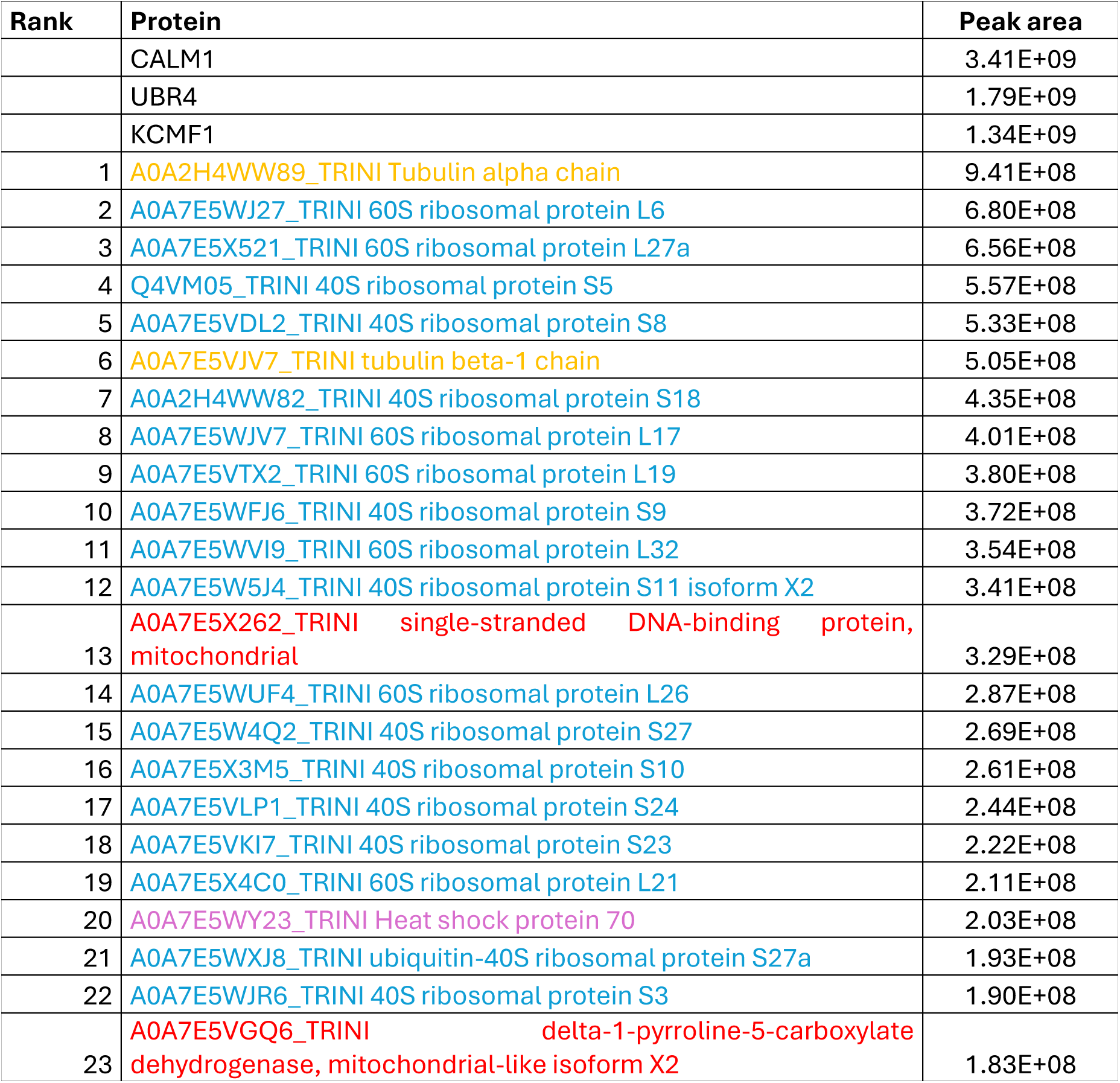
MS analysis of *Trichoplusia ni* contaminants in the purified human UBR4 complex. Mitochondrial proteins are colored in red, while common contaminants are colored in blue (ribosome), orange (tubulin) and purple (chaperones).

**Table S3.**
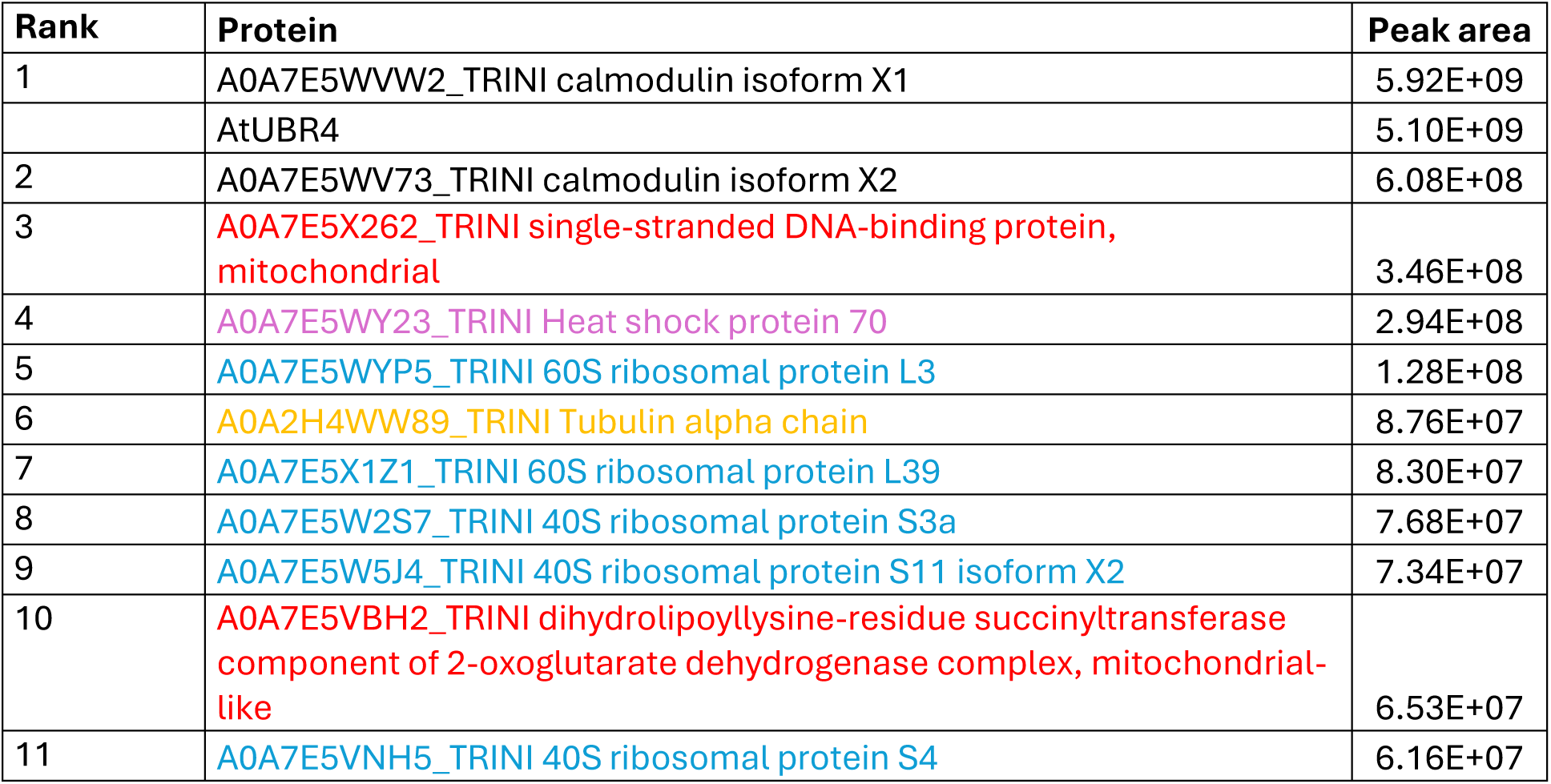
MS analysis of *Trichoplusia ni* contaminants in purified AtUBR4. Mitochondrial proteins are colored in red, while common contaminants are colored in blue (ribosome), orange (tubulin) and purple (chaperones).

**Table S4.** MS analysis of *Trichoplusia ni* contaminants in the purified CeUBR4 complex. Mitochondrial proteins are colored in red, while common contaminants are colored in blue (ribosome), orange (tubulin) and purple (chaperones).

| Rank | Protein | Peak area |
| --- | --- | --- |
|  | KCMF1 | 7.40E+09 |
|  | UBR4 | 4.77E+09 |
| 1 | A0A2H4WW89_TRINI Tubulin alpha chain | 1.19E+09 |
| 2 | A0A7E5VJV7_TRINI tubulin beta-1 chain | 7.49E+08 |
| 3 | A0A7E5X521_TRINI 60S ribosomal protein L27a | 5.01E+08 |
| 4 | A0A7E5WXJ8_TRINI ubiquitin-40S ribosomal protein S27a | 4.77E+08 |
| 5 | A0A7E5WJQ8_TRINI 40S ribosomal protein S6 | 4.76E+08 |
| 6 | A0A7E5WFJ6_TRINI 40S ribosomal protein S9 | 3.99E+08 |
| 7 | A0A7E5VDL2_TRINI 40S ribosomal protein S8 | 3.40E+08 |
| 8 | A0A7E5WBC0_TRINI 40S ribosomal protein S14 | 3.23E+08 |
| 9 | A0A7E5WYR8_TRINI 40S ribosomal protein S2 | 2.80E+08 |
| 10 | A0A7E5WY23_TRINI Heat shock protein 70 | 2.75E+08 |
| 11 | A0A7E5WJR6_TRINI 40S ribosomal protein S3 | 2.65E+08 |
| 12 | A0A7E5W5J4_TRINI 40S ribosomal protein S11 isoform X2 | 2.59E+08 |
| 13 | A0A7E5WJV7_TRINI 60S ribosomal protein L17 | 2.53E+08 |
| 14 | A0A7E5W4Q2_TRINI 40S ribosomal protein S27 | 2.46E+08 |
| 15 | A0A7E5W8Z9_TRINI 40S ribosomal protein S16 isoform X1 | 2.13E+08 |
| 16 | A0A2H4WW82_TRINI 40S ribosomal protein S18 | 1.85E+08 |
| 17 | A0A7E5VWI8_TRINI eukaryotic translation initiation factor 3 subunit K | 1.74E+08 |
| 18 | A0A7E5WBC6_TRINI 60S ribosomal protein L24 | 1.66E+08 |
| 19 | Q4VM05_TRINI 40S ribosomal protein S5 | 1.51E+08 |
| 20 | A0A7E5W2S7_TRINI 40S ribosomal protein S3a | 1.36E+08 |
| 21 | A0A7E5WVI9_TRINI 60S ribosomal protein L32 | 1.36E+08 |
| 22 | A0A7E5X5U8_TRINI 60S ribosomal protein L4 | 1.26E+08 |
| 23 | A0A7E5VTX2_TRINI 60S ribosomal protein L19 | 1.20E+08 |
| 24 | A0A7E5X3M5_TRINI 40S ribosomal protein S10 | 1.15E+08 |
| 25 | A0A7E5WLS0_TRINI 40S ribosomal protein S26 | 1.11E+08 |
| 26 | A0A7E5VNH5_TRINI 40S ribosomal protein S4 | 9.46E+07 |
| 27 | A0A7E5X4C0_TRINI 60S ribosomal protein L21 | 8.94E+07 |
| 28 | A0A7E5VYY3_TRINI 60S ribosomal protein L9 | 7.95E+07 |
| 29 | A0A7G9U7K2_TRINI Heat shock protein 21.4 | 7.53E+07 |
| 30 | A0A7E5W0B5_TRINI 60S ribosomal protein L18a | 6.65E+07 |
| 31 | A0A7E5X019_TRINI 60S ribosomal protein L13a | 5.67E+07 |
| 32 | A0A7E5VKI7_TRINI 40S ribosomal protein S23 | 5.39E+07 |
| 33 | A0A7E5W784_TRINI 40S ribosomal protein S13 | 4.65E+07 |
| 34 | A0A7E5WY71_TRINI 40S ribosomal protein S15Aa | 4.33E+07 |
| 35 | A0A7E5WEG5_TRINI 60S ribosomal protein L35 | 4.27E+07 |
| 36 | A0A7E5WQP8_TRINI 60S ribosomal protein L10 | 4.13E+07 |
| 37 | A0A7E5VLP1_TRINI 40S ribosomal protein S24 | 3.81E+07 |
| 38 | A0A7E5X2F1_TRINI probable medium-chain specific acyl-CoA dehydrogenase, mitochondrial isoform X3 | 3.79E+07 |
| 39 | A0A7E5X5Y1_TRINI tubulin beta chain-like | 3.34E+07 |
| 40 | A0A7E5WJ27_TRINI 60S ribosomal protein L6 | 3.32E+07 |
| 41 | A0A7E5VE68_TRINI 60S ribosomal protein L35a | 3.17E+07 |
| 42 | A0A7E5WAV9_TRINI dnaJ homolog subfamily A member 2-like | 3.13E+07 |
| 43 | A0A7E5WF28_TRINI 60S ribosomal protein L27-like | 2.83E+07 |
| 44 | A0A7E5VSK5_TRINI heat shock protein 83 | 2.79E+07 |
| 45 | A0A7E5VV98_TRINI 60S ribosomal protein L13 | 2.77E+07 |
| 46 | A0A7E5X1Z1_TRINI 60S ribosomal protein L39 | 2.77E+07 |
| 47 | A0A7E5WUF4_TRINI 60S ribosomal protein L26 | 2.60E+07 |
| 48 | A0A7E5WME8_TRINI 60S ribosomal protein L34-like | 2.52E+07 |
| 49 | A0A7E5X262_TRINI single-stranded DNA-binding protein, mitochondrial | 2.17E+07 |
| 50 | A0A7E5WNQ1_TRINI 60S ribosomal protein L23 | 1.97E+07 |

